# Incorporating exon-exon junction reads enhances differential splicing detection

**DOI:** 10.1101/2025.01.29.631417

**Authors:** Mai T. Pham, Michael J.G. Milevskiy, Jane E. Visvader, Yunshun Chen

**Affiliations:** ACRF Cancer Biology and Stem Cells Division, The Walter and Eliza Hall Institute of Medical Research, Parkville, VIC 3052, Australia; Bioinformatics Division, The Walter and Eliza Hall Institute of Medical Research, Parkville, VIC 3052, Australia; Department of Medical Biology, The University of Melbourne, Parkville, VIC 3010, Australia

## Abstract

**Motivation:** RNA sequencing (RNA-seq) is a gold standard technology for studying gene and transcript expression. Different transcripts from the same gene are usually determined by varying combinations of exons within the gene, formed by splicing events. One method of studying differential alternative splicing between groups in short-read RNA-seq experiments is through differential exon usage (DEU) analyses, which use exon-level read counts along with downstream statistical testing strategies. Popular pipelines for these analyses include *Rsubread* for read alignment and exon-level count quantification, and *edgeR* and *limma* for statistical testing of differential splicing. However, the standard *Rsubread::featureCounts* count summarization method does not consider exon-junction information, which may reduce the statistical power in differential splicing analyses.

**Results:** A new feature quantification approach is proposed to incorporate both exon and exon-junction reads into the popular *Rsubread-edgeR* and *Rsubread-limma* frameworks for detecting differential splicing events. This new Differential Exon-Junction Usage (DEJU) analysis pipeline demonstrates increased statistical power compared to existing popular methods while effectively controlling the false discovery rate.

**Availability and implementation:** Data and code to reproduce the results shown in this article are available from https://github.com/TamPham271299/DEJU.

## Introduction

Alternative splicing is a post-transcriptional regulatory mechanism that occurs in approximately 95% of multi-exon genes in higher eukaryotes (Pan *et al*., 2008). This process enables a single gene to produce various mRNA isoforms, thereby significantly enhancing the transcriptome and protein diversity to meet the functional demands of different cell types (Graveley, 2001). Furthermore, alternative splicing plays a crucial, yet not fully understood, role in carcinogenesis. Aberrant expression of abnormal isoforms of cancer-related genes due to alternative splicing can contribute to cancer progression, metastasis, and drug resistance (Zhang and Manley, 2013; Wang and Lee, 2018; Oh *et al*., 2021).

Over the past decade, bulk RNA-sequencing (RNA-seq) has emerged as a powerful technology for measuring the expression levels of genomic features such as genes, transcripts, and exons. With the decreasing cost of sequencing, achieving higher sequencing depths has become both feasible and beneficial for detecting differential splicing between conditions. Consequently, many computational methods and software tools have been developed to utilize bulk RNA-seq data for more accurate and efficient differential splicing detection (Mehmood *et al*., 2020).

In a short-read bulk RNA-seq experiment, an effective method to detect alternative splicing events is by testing for differential exon usage (DEU). This approach enables the rapid identification of genes that may be differentially spliced, referred to hereafter as DEU genes, without requiring the specification of the exact isoforms present in different groups. For example, the *featureCounts* function in the *Rsubread* package can quantify read counts at the exon level (Liao *et al*., 2014). These exon-level count data can then be utilized for DEU analysis using the *diffSpliceDGE* function in the *edgeR* package (Chen *et al*., 2024) or the *diffSplice* function in the *limma* package (Ritchie *et al*., 2015).

Exon-level reads can be broadly classified into two categories: internal exon reads, which map entirely within exons, and exon-exon junction reads, which span across two exons. In the current implementation of *featureCounts* in the *Rsubread* package, if a read maps to a region spanning two exons, it will, by default, contribute one count to both exons, leading to a double-counting issue. Moreover, this current approach was originally developed to identify differential usage of flattened and merged exons, but it is limited in its ability to detect exon extension or truncation involving alternative splice sites, retained introns, or nested exon skipping. This limitation arises from the lack of information regarding junction sites.

To address the double-counting problem of exon-level read counts and enhance the overall performance of DEU analysis within the *Rsubread-edgeR-limma* framework, we propose a differential exon-junction usage (DEJU) analysis pipeline. This pipeline incorporates exon-junction reads into the existing DEU analysis process. Specifically, we utilize splice-aware aligners such as STAR (Dobin *et al*., 2013) to identify splice junctions and subsequently quantify junction reads using the *featureCounts* function with settings that differentiate internal exon reads from exon-junction reads. In the final read count matrix, exon-junctions are included as features alongside exons. This approach ensures that each exon-junction read is uniquely assigned to a single feature, thereby effectively resolving the double-counting issue and ensuring that the resulting library sizes accurately reflect the true number of sequence reads.

In this article, we benchmark the performance of our proposed DEJU pipeline against the current DEU approaches and other popular DEU analysis methods such as *DEXseq* (Anders *et al*., 2012) and *Junction-Seq* (Hartley and Mullikin, 2016). We use simulated bulk RNA-seq datasets with various designed splicing patterns, where the ground truths are known, to evaluate the accuracy and effectiveness of these methods. The DEJU pipeline is also evaluated for its robustness in terms of precision and sensitivity in detecting DEU genes across various combinations of library sizes and sample sizes. We show that our DEJU analysis pipeline is more precise, powerful, and flexible in detecting a broader range of splicing events compared to existing popular methods. It enhances statistical power while effectively controlling the false discovery rate. The DEJU analysis process is also efficient in time and memory usage. Additionally, this pipeline was tested on a mouse bulk RNA-seq experiment involving two mammary epithelial cell (MEC) types (luminal progenitor and mature luminal) (Milevskiy et al., 2023), revealing biologically meaningful findings related to alternative splicing that could not be detected previously.

## Methods

### Reference genome

We retrieved the primary genome sequence FASTA file and the comprehensive gene annotation GTF file for the GRCm39 mouse genome release M32 from the GENCODE database (https://www.gencodegenes.org). The GTF file was used to generate comprehensive annotations for genomic features, including flattened exons and splice junctions. To create flattened and merged exon annotation, the *flattenGTF* function of the *Rsubread* package (Liao *et al*., 2019) was employed to extract exonic regions from the GTF file and merge overlapping exons attributed to the same gene. In addition, a splice junction annotation file, referred to hereafter as junction database, was generated from annotated exonic regions at the transcript level. Two resultant annotation files were saved in a standard annotation format (SAF). See Supplementary Methods section 3.1 for more details.

### Simulated datasets

#### Customized transcriptome

To create a customized transcriptome for simulation studies, a subset of 5000 multi-exon protein-coding genes with two transcripts were randomly generated. The annotated exonic regions of the selected genes were extracted and the overlapping exons attributed to the same gene were flattened and merged. Transcripts from the resulting transcriptome were manually modified to mimic the common alternative splicing patterns: exon skipping (ES), mutually exclusive exon (MXE), alternative 3’/5’ splice site (ASS), and intron retention (IR) in equal proportions. See Supplementary Methods section 3.2 for more details.

#### Simulation of RNA-seq sequence reads

The *simReads* function in the Bioconductor package *Rsubread* was used to simulate 75 base pair (bp) long paired-end RNA-seq sequence reads based on the customized transcriptome and the outputs were stored in FASTQ format. The baseline expression levels of transcripts were simulated following the Zipf’s law (Furusawa and Kaneko, 2003). Biological variations between replicates within each group were simulated following a gamma distribution. Different library sizes were considered in simulation. In a balanced scenario, the library size was set at 50 million reads across all the samples, whereas in an unbalanced case they were alternating between 25 million and 100 million reads over all samples. Two groups of sample were simulated with different number of biological replicates per group (3, 5, or 10) under different scenarios. To simulate genuine differential splicing events, 1000 out of the total 5000 genes were randomly selected and the four common alternative splicing patterns (ES, MXE, ASS, and IR) were simulated for those 1000 selected genes in equal proportions. For each of those 1000 genes, the baseline abundance of one isoform was increased in the first group and the other isoform in the second group multiplicatively by a fold-change of 3. Null simulations in which underlying transcript abundances are consistent across replicates of both groups were also generated to assess type I error rate control. For each combination of library size and number of biological replicates per group, the simulation was run 20 times, and the results represent the average across those 20 runs. See Supplementary Methods section 3.2 for more details.

### Differential exon-junction usage analysis workflow

We developed a complete workflow for differential exon-junction usage (DEJU) analysis of short-read bulk RNA-sequencing experiments. This workflow utilizes the *STAR* and *Rsubread* for read alignment and feature quantification respectively, and it leverages the *edgeR* or *limma* packages for downstream statistical analysis (Figure 1). Details of key workflow steps are described below.

**Figure 1:**
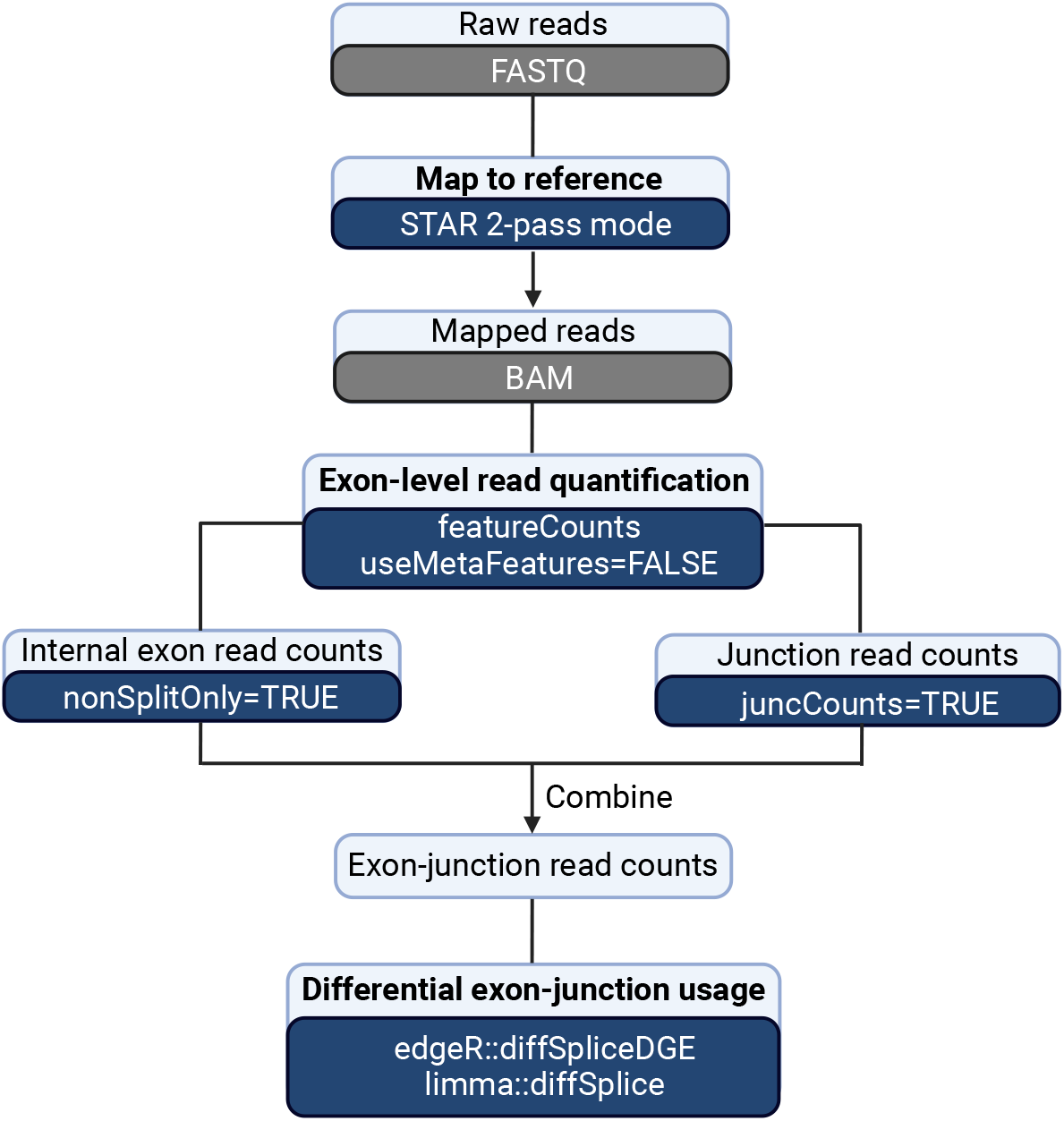
The Rsubread-edgeR-limma framework of the DEJU analysis pipeline.

#### Data pre-processing

Simulated RNA-seq reads were aligned to the mouse reference genome GRCm39 using *STAR* aligner using the 2-pass mapping mode with re-generated genome. To achieve the highest sensitivity to novel junction detection, a list of junctions detected by the 1-pass mapping from all samples of compared conditions were collapsed and filtered. Any junctions supported by fewer than three uniquely mapping reads across all samples were removed from the analysis. The resulting set of junctions were subsequently used to reindex the reference genome for the second round of mapping. Only junction reads aligned to junctions that passed filtering were kept in the alignment BAM files using the `--outFilterType BySJout` option.

#### Exon and junction read quantification

The aligned reads were then quantified by the *featureCounts* function using the flattened and merged exon annotation with the argument `useMetaFeatures=FALSE`. The `nonSplitOnly` argument in *featureCounts* allows quantification of internal exon read counts, whereas the `juncCounts` argument enables *featureCounts* to return an extra count matrix for exon-exon junctions. By default, both `nonSplitOnly` and `juncCounts` are set to `FALSE`. Under the DEJU pipeline, the read quantification step was performed separately using two different settings, `nonSplitOnly=TRUE` and `juncCounts=TRUE`, to get internal exon read counts and junction read counts, respectively. The reference genome-generated junction database was subsequently incorporated into our DEJU pipeline to improve the assignment of annotated junctions to genes. Internal exon and processed junction read counts were then combined into one exon-junction read count matrix. Exons and junctions with low number of mapped reads were filtered by the *filterByExpr* function in *edgeR*. The trimmed mean of M values (TMM) normalization (Robinson and Oshlack, 2010) was performed using the *normLibSizes* function in the *edgeR* package to account for the composition biases between libraries.

#### Differential exon-junction usage detection

The downstream differential exon-junction usage analyses were performed using the *diffSpliceDGE* function in *edgeR* or the *diffSplice* function in *limma*. These two functions identified features (both exons and junctions) that were used differentially between the groups. The feature-level test results were then summarized at the gene level using either the Simes method (Simes, 1986), which combines feature-level p-values within each gene, or an *F*-test, which combines feature-level quasi-likelihood *F*-test statistics for that gene (Chen *et al*., 2024). Under both the Simes method and the *F*-test, genes were considered to exhibit differential exon-junction usage if the gene-level false discovery rate (Benjamini and Hochberg, 1995) is below 0.05.

### Methods benchmarking and evaluation

We assessed the performance of the DEJU pipeline and other popular approaches in terms of their statistical power in detecting DEU genes, the control of FDR, as well as their computational resource usage. In particular, the *diffSpliceDGE* function of *edgeR* and the *diffSplice* function of *limma* were used for downstream statistical testing as part of the DEJU pipeline, denoted as *DEJU-edgeR* and *DEJU-limma*, respectively. These two functions were also used in the standard DEU pipelines, denoted as *DEU-edgeR* and *DEU-limma*, where the RNA sequencing reads were assigned to exons only using the default settings of *featureCounts* in *Rsubread*. Other pipelines being considered in this benchmarking study include *DEXSeq* and *JunctionSeq*. We used the *featureCounts* function in *subread* package to preprocess exon counts for DEXSeq, which was followed by the DEU analysis using the *DEXSeq* function in *DEXSeq* package with default settings. FDR at the gene-level was controlled using the *perGeneQValue* function of *DEXSeq* package. On the other hand, we used *QoRTs* (Hartley and Mullikin, 2015) software package with `--runFunctions writeKnownSplices,writeNovelSplices,writeSpliceExon` *options to generate exon and splice junction counts which were in turn used as input of JunctionSeq* to test for differentials of exon and splice junctions using default settings.

To evaluate the robustness and accuracy in the DEU analysis of the compared tools, different simulation settings were applied, considering the number of samples per group and balanced/unbalanced library size scenarios. The maximum memory usage and the computing speed of the DEU/DEJU analysis were also recorded for each compared pipeline.

Full details of the simulation and benchmarking procedures are included in Supplementary Methods, Section 3.2.

### RNA-seq profiles of mouse mammary epithelium

All benchmarked pipelines in the simulation study were applied to an Illumina short-read paired-end RNA-seq experiment exploring the adult mouse epithelial mammary gland (Milevskiy *et al*., 2023). Three biological replicate samples were obtained for each of the two cell populations of interest, including luminal progenitor (LP) and mature luminal (ML). The raw sequencing data in FASTQ format of all six samples were downloaded from the NCBI Gene Expression Omnibus (GEO) series GSE227748. Paired-end reads were first mapped to the mouse reference genome using the *STAR* aligner with the 2-pass mode. The same reference genome set was used for the DEU/DEJU analysis of the case study. To improve the overall mapping accuracy, non-canonical junctions were excluded using an extra option `--outFilterIntronMotifs RemoveNoncanonical` in the second pass. Genes without valid gene symbols were also removed from the analysis. Filtering, normalization, and downstream statistical testing were conducted using the same methods for each pipeline as described in the previous simulation study. Full details of the case-study analyses are included in Supplementary Methods section 3.3.

## Results

### False discovery rate and statistical power

We evaluated the performance of various DEU and DEJU pipelines with respect to their power in detecting DEU genes featuring different splicing patterns and their ability to control FDR. See Supplementary Table S1 for full results. Table 1 showed the statistical power and FDR of *DEJU-edgeR* and *DEJU-limma* on simulated datasets under different simulation settings and for different splicing patterns. In general, the ability to detect DEU genes increased with larger sample sizes, particularly evident in the improved detection of ASS and IR events. *DEJU-edgeR* effectively controlled FDR at the nominal rate of 0.05 for all splicing events, although it was slightly more conservative compared to *DEJU-limma*. In contrast, *DEJU-limma* struggled to control the FDR as effectively as *DEJU-edgeR*, particularly in detecting MXE events.

**Table 1:**
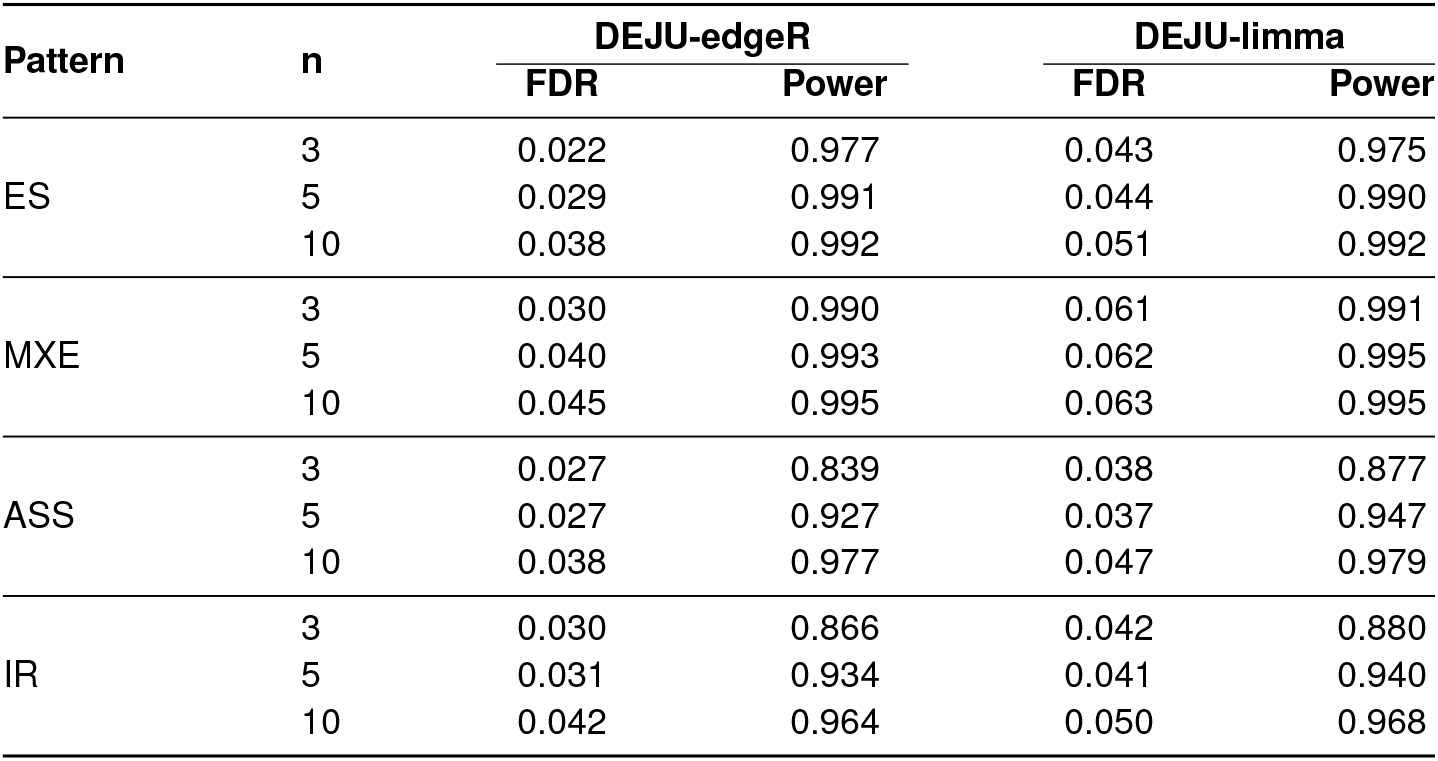
DEU detection FDR and power of *DEJU-edgeR* and *DEJU-limma* using the gene-level Simes method under different sample sizes (n). The table shows the FDR and power in DEU detection for simulations with balanced library sizes and 250 genuine DEU genes for each splicing pattern. Results were averaged over 20 independent simulation runs for each sample size.

| Pattern | n | DEJU-edgeR |  | DEJU-limma |  |
| --- | --- | --- | --- | --- | --- |
|  |  | FDR | Power | FDR | Power |
| ES | 3 | 0.022 | 0.977 | 0.043 | 0.975 |
|  | 5 | 0.029 | 0.991 | 0.044 | 0.990 |
|  | 10 | 0.038 | 0.992 | 0.051 | 0.992 |
| MXE | 3 | 0.030 | 0.990 | 0.061 | 0.991 |
|  | 5 | 0.040 | 0.993 | 0.062 | 0.995 |
|  | 10 | 0.045 | 0.995 | 0.063 | 0.995 |
| ASS | 3 | 0.027 | 0.839 | 0.038 | 0.877 |
|  | 5 | 0.027 | 0.927 | 0.037 | 0.947 |
|  | 10 | 0.038 | 0.977 | 0.047 | 0.979 |
| IR | 3 | 0.030 | 0.866 | 0.042 | 0.880 |
|  | 5 | 0.031 | 0.934 | 0.041 | 0.940 |
|  | 10 | 0.042 | 0.964 | 0.050 | 0.968 |

Figure 2 showed the observed number of true positive and false positive DEU genes for all 6 benchmarked pipelines including *DEJU-edgeR, DEJU-limma, DEU-edgeR, DEU-limma, DEXSeq*, and *JunctionSeq* under a nominal FDR control of 0.05, for the simulation scenario with three samples per group and balanced library size. Full details on FDR control across other simulation scenarios are provided in Supplementary Figure S1.

**Figure 2:**
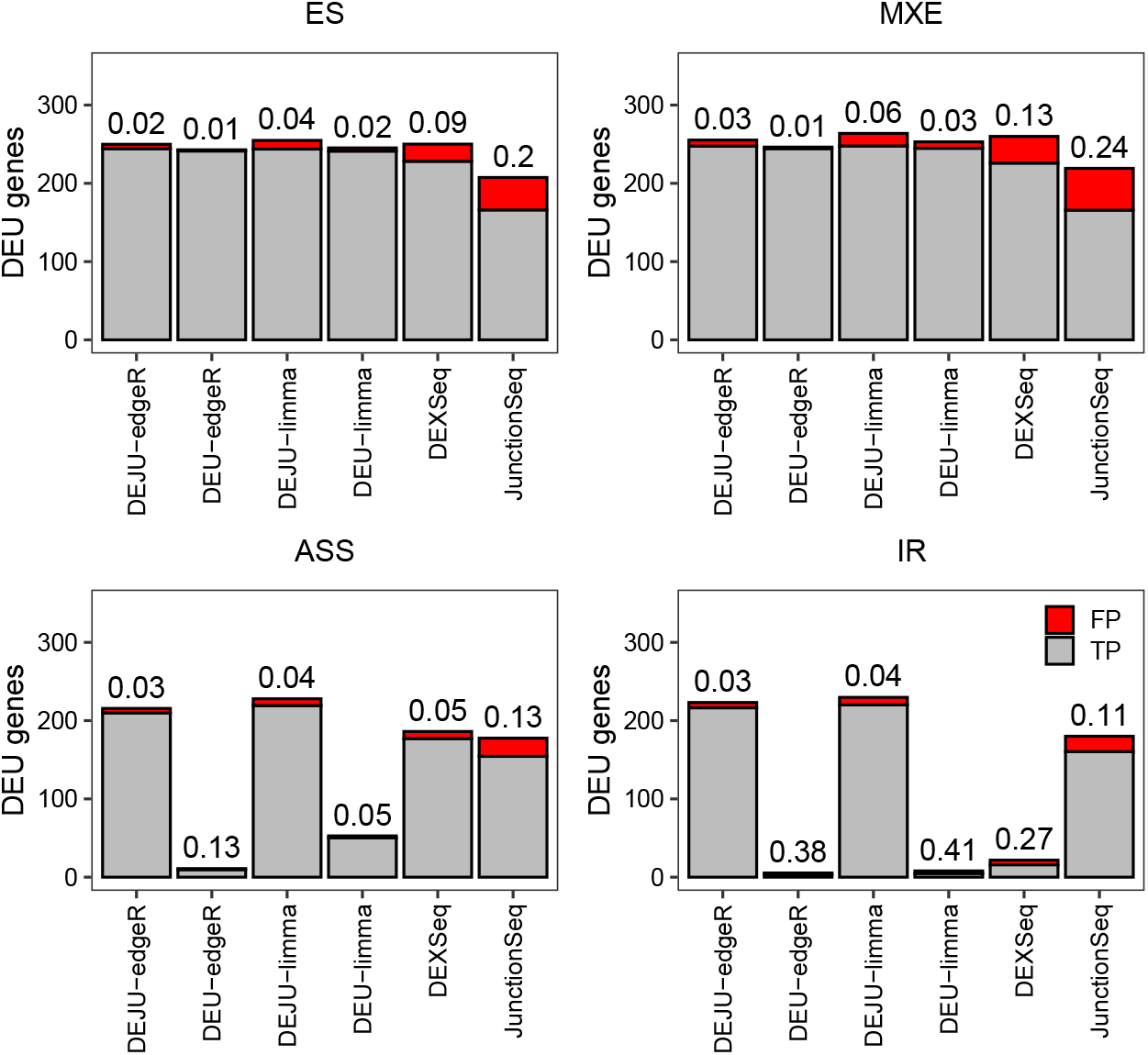
FDR of 6 benchmarked pipelines in the DEU analysis. Stacked barplots showing the average number of true (gray) and false (red) positive DEU genes at nominal 5% FDR in different splicing pattern scenarios – ES, MXE, ASS, and IR. The observed FDR is shown over each bar. Results are averaged over 20 independent simulation runs with 75 bp paired-end reads and 250 genuine DEU genes simulated for each splicing pattern. DEU/DEJU analysis results are shown using the gene-level Simes method.

To further evaluate methods for FDR control, we assessed the number of false discoveries in the set of top-ranked most significant DEU genes identified by each method (Figure 3). Overall, *DEJU-edgeR* and *DEJU-limma* consistently produced the smallest number of false discoveries among all methods for any number of top-ranked DEU genes. Across various library and sample size configurations, *DEU-edgeR, DEU-limma, DEXSeq*, and *JunctionSeq* exhibited more false positive DEU genes for any given number of top-ranked DEU genes. However, increasing the number of replicates per group led to a slight reduction in false discoveries for most benchmarked methods. Compared to the gene-level Simes method, DEU and DEJU analyses using the gene-level *F*-test resulted in slightly higher numbers of false discoveries. Nevertheless, these methods produced comparable results in scenarios with 10 replicates per group (Supplementary Figure S2).

**Figure 3:**
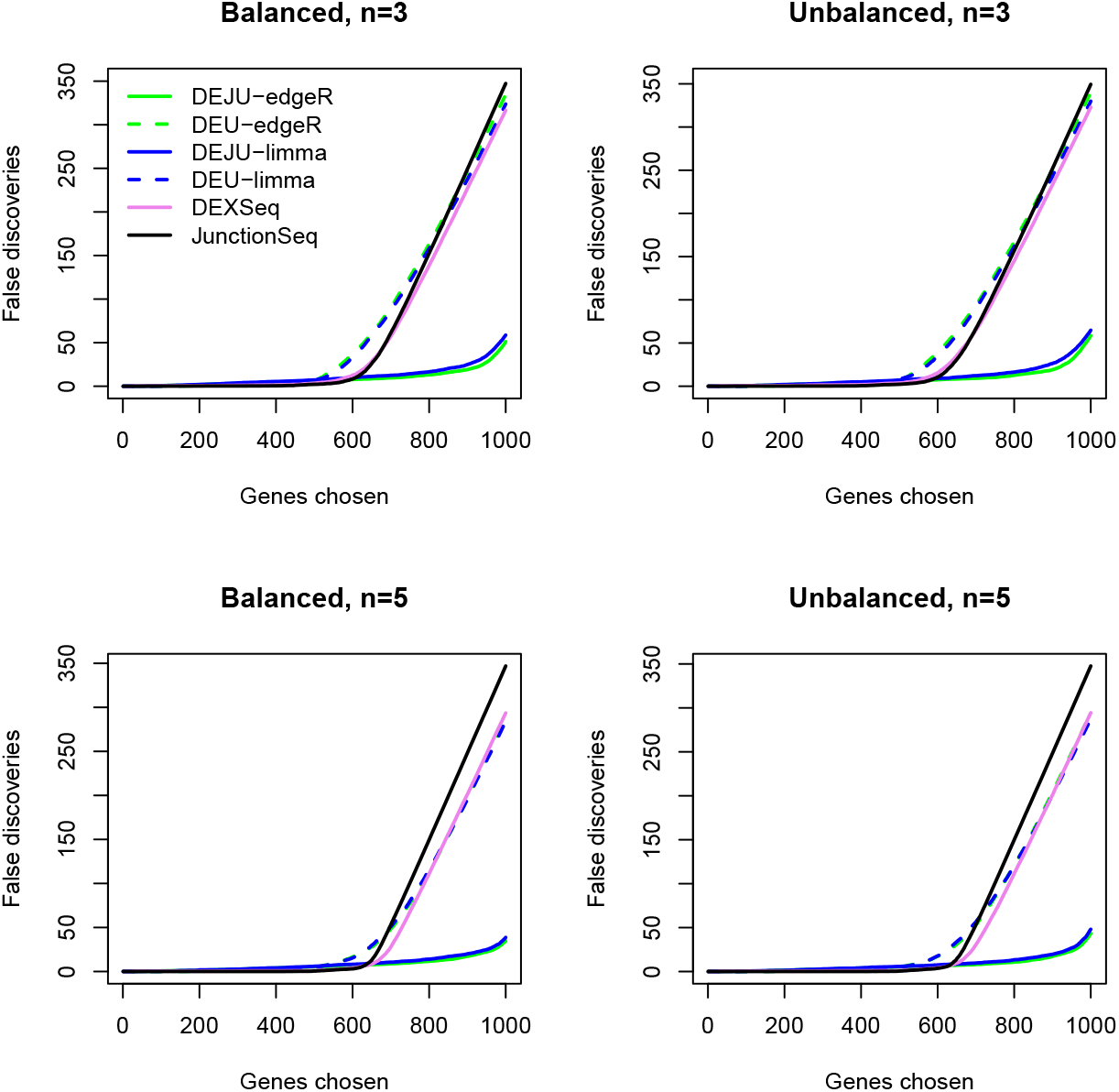
Panels show the average number of false discoveries as a function of the number of chosen DEU genes across the combination of balanced/unbalanced library size and sample sizes (3,5 replicates per group). Results are averaged over 20 independent simulation runs with 75 bp paired-end reads. DEU/DEJU analysis results are shown using the gene-level Simes method.

The two DEJU pipelines consistently gave an empirical FDR below the nominal FDR of 0.05 and showed a substantially higher power and flexibility in detecting any of the designed splicing patterns compared to other methods, regardless of the number of replicates per group and library sizes (Supplementary Figures S3-S4).

### Computational resources

We compared the computational speed and resources required for the downstream statistical testing part under different pipelines. Both *DEU-edgeR* and *DEJU-edgeR* uses the *diffSpliceDGE* function for statistical analysis, whereas both *DEU-limma* and *DEJU-limma* uses the *diffSplice*. Table 2 summarized runtime and memory cost for each method in analyzing RNA-seq reads with the balanced library sizes across samples, benchmarked in both parallel and serial modes to illustrate work-load distribution and method efficiency. Based on these metrics, we observed that both *limma* and *edgeR* pipelines, regardless of DEU or DEJU, had substantially lower compute cost and turnaround time, while consistently maintaining their accuracy and power in detecting DEU genes with different splicing patterns. Additionally, the computational time and resources of *DEXSeq* and *JunctionSeq* scaled up proportionally with increasing sample sizes, further emphasizing the efficiency of *limma* and *edgeR* pipelines in DEU analysis. The consumption in computing resources of these methods for unbalanced library sizes is shown in Supplementary Table S2.

**Table 2:** Compute cost of compared DEU detection tools with increasing sample sizes (n). All tests are carried out on Intel Xeon E5-2690 v4 server with data on local drive. Parallel tests use 4 cores, serial tests use 1 core. Memory columns show peak RSS. Walltime and memory usage of the *diffSpliceDGE* function of *edgeR* and the *diffSplice* function of *limma* is for both DEU and DEJU analyses. No parallelization is available for these methods.

| n | Method | Walltime (min) |  | Memory (Gb) |  |
| --- | --- | --- | --- | --- | --- |
|  |  | Parallel | Serial | Parallel | Serial |
| 3 | edgeR | - | 0.47 | - | 0.64 |
|  | limma | - | 0.33 | - | 0.55 |
|  | DEXSeq | 9.74 | 18.29 | 4.81 | 2.71 |
|  | JunctionSeq | 10.12 | 16.06 | 6.87 | 3.11 |
| 5 | edgeR | - | 0.64 | - | 0.63 |
|  | limma | - | 0.42 | - | 0.63 |
|  | DEXSeq | 11.56 | 32.85 | 4.91 | 3.37 |
|  | JunctionSeq | 10.57 | 24.54 | 8.61 | 4.24 |
| 10 | edgeR | - | 0.71 | - | 0.80 |
|  | limma | - | 0.41 | - | 0.75 |
|  | DEXSeq | 29.15 | 92.55 | 6.27 | 4.99 |
|  | JunctionSeq | 20.55 | 58.39 | 11.36 | 5.34 |

### Differential exon-junction usage analysis for mouse mammary epithelial cell lines

Here, we detected DEU genes between LP and ML MECs from the Illumina short paired-end read RNA-seq experiments with three biological replicate samples per cell line. Libraries were sequenced with an Illumina NextSeq 500 sequencing system, generating 34-82 millions of 80-81 bp read-pairs per sample. DEU genes were identified using our DEJU analysis pipeline, and the library sizes of internal exon counts, junction counts, and exon-junction counts of *DEU-edgeR* and *DEJU-edgeR* were recorded (Supplementary Table S3). Overall, the proposed DEJU pipeline gave library sizes more precisely - 37-86 million reads - as the actual sequencing libraries compared to the existing DEU analysis pipeline. Diagnostic plots of the exon-junction read quantification of DEU/DEJU analysis pipeline implemented in *edgeR* are shown in Supplementary Figure S5.

Figure 4 showed results of DEU gene detection between LP and ML MECs by *DEU-edgeR* and *DEJU-edgeR*. Overall, about 55% of positive discoveries were exclusively reported by *DEJU-edgeR* using the gene-level Simes and *F*-test (Figure 4A). Full lists of DEU genes only detected by *DEJU-edgeR* with the gene-level Simes method or *F*-test can be accessed via Supplementary Table S4.

**Figure 4:**
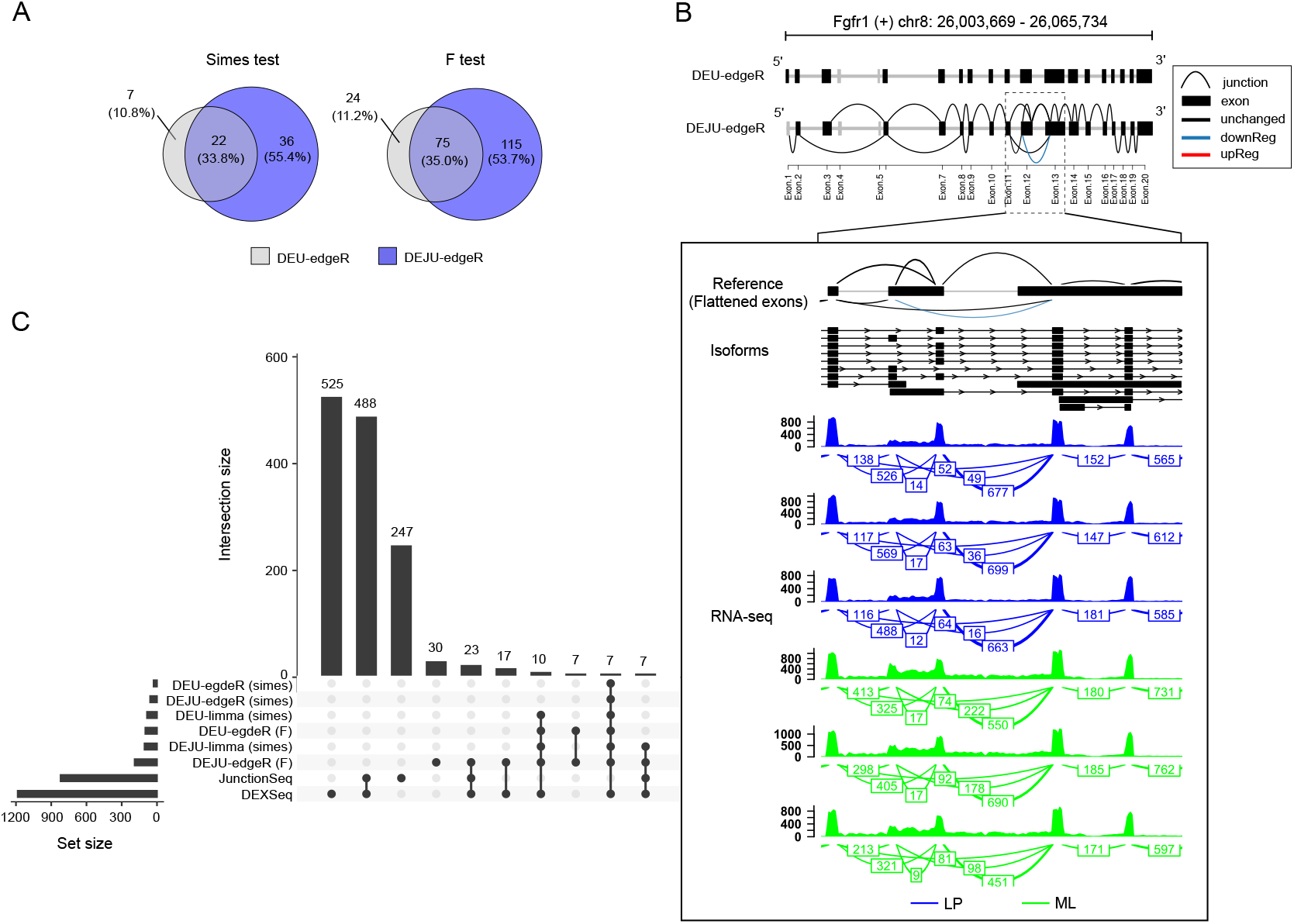
Panels (A)-(C) shows DEU detection results between LP and ML samples. (A) VennDiagram shows the number of DEU gene detections by *DEU-edgeR* (gray) and *DEJU-edgeR* (purple); (B) An example of a DEU gene, *Fgfr1*, with a down-regulated junction in LP samples exclusively detected by *DEJU-edgeR* with the gene-level Simes method (adjusted *P*-value = 0.018) compared to *DEU-edgeR* (adjusted *P*-value = 1). (Top panels) Schematic exon-junction plots showing the up-regulated (red), down-regulated (blue), and non-differentially (black) used exons and junctions in LP against ML samples; (Middle panels) a UCSC comprehensive transcript annotation track of mm39; (Bottom panels) Sashimi plots alongside the coverage of RNA-seq reads of LP (blue) and ML (green) samples. Exon-exon junction read counts are shown in a box; (C) Upset plots showing the intersection of different DEU gene sets detected by bench-marked pipelines.

Of particular interest was the detection of *Fgfr1, Myl6*, and *Mvb12a*, whose roles have been reported in signaling pathways crucial for MEC differentiation and development. For instance, Zhao *et al*. (2019) observed the differential expression of spliced variants of *Fgfr1* that govern cell differentiation. Similarly, the expression levels of *Myl6* have been identified as a potential marker for distinguishing epithelial cell populations (Sleeman *et al*., 2005; Li *et al*., 2020). Furthermore, *Mvb12a* has been implicated in the regulation of the epidermal growth factor receptor (EGFR) signaling pathway (Morato *et al*., 2021). While alternative splicing events involving these genes have been modestly reported in previous studies, the splicing signals identified in our study warrant further investigation to elucidate their potential roles in regulating MEC differentiation and development.

Figure 4B highlights the differentially used junctions of *Fgfr1* detected by *DEJU-edgeR* (adjusted *P*-value = 0.018) but not by *DEU-edgeR* (adjusted *P*-value = 1). See Supplementary Figure S6 for more illustrative examples of DEU genes exclusively detected by *DEJU-edgeR* with the gene-level Simes method or *F*-test. Figure 4C visualizes the intersections among different sets of DEU genes detected by benchmarked methods. Overall, the DEJU analysis pipeline implemented in *edgeR* identified double the number of positive discoveries compared to the DEU pipeline (Supplementary Table S5). Of DEU/DEJU analysis pipelines, *DEJU-edgeR* with the gene-level *F*-test detected the highest number of DEU genes. Only a minority of DEU genes were detected by all benchmarked methods. On the other hand, a substantial number of positive discoveries exclusively detected by *DEXSeq* and *JunctionSeq*. Full intersection sets are shown in Supplementary Figure S7. More examples of DEU genes detected by *DEJU-limma, DEXSeq*, and *JunctionSeq* are further visualized in Supplementary Figures S8-10.

## Discussion and Conclusion

Here, we present a differential exon-junction usage (DEJU) analysis pipeline that enables fast and accurate identification of genes exhibiting differential splicing events between experimental conditions. This DEJU pipeline enhances read mapping accuracy by utilizing the two-pass mapping strategy implemented in STAR. It separates exon-junction reads from internal exon reads and incorporates both types of reads in the downstream analysis. The new *featureCounts* read-counting strategy adopted in the DEJU pipeline resolves the double-counting issue that currently exists in standard DEU pipelines. Furthermore, our DEJU pipeline proves advantageous in identifying a greater number of DEU genes, especially ones with alternative splice sites and retained introns, as it captures signals from both exons and junctions, leading to more comprehensive detection. Additionally, it allows for the reassignment of junctions to genes using junction databases as references, further improving the accuracy of junction read counts. The downstream statistical testing of the DEJU pipeline is performed using either the *diffSpliceDGE* function in *edgeR*, or the *diffSplice* function in *limma*, both of which have been shown to be highly effective and computational efficient.

We perform comprehensive simulation studies to benchmark the performance of the DEJU pipeline with many other existing popular DEU methods. Our simulation results highlight the superior performance of the DEJU approach in detecting different splicing patterns, especially ASS and IR, with regard to its statistical power and its ability to control FDR compared to other DEU analysis strategies. Our case study of the mouse mammary gland highlights the benefits of the DEJU pipeline. It reveals genes previously not identified as DEU genes, and these newly identified genes have meaningful biological interpretations.

Although the DEJU pipeline has been proven to outperform other counterparts notably, it can be optimized to some extent. For example, we can incorporate the exon binning strategy applied in *DEXSeq* and *JunctionSeq* into the exon-junction count summarization. One suggestion is to quantify internal exon reads to exon counting bins, instead of merged exons, while summarizing junction read counts separately, thus would enhance the sensitivity in the DEU detection to a certain extent. However, the statistical power and the ability to control the FDR of the modified DEJU pipeline need to be examined if implemented.

As an efficient and powerful method to detect DEU genes, our proposed DEJU analysis workflow could be applied to further explore the other differential splicing studies such as those in cancer research. In particular, we can explore RNA-seq data from various cancers in the TCGA public database to detect novel splicing events within cancer-related genes or identify splicing events associated with drug resistance, suggesting potential targets for further biological and clinical cancer research.

## Author contributions statement

M.T.P. and Y.C. conceived the experiment and conducted the simulation studies. M.T.P. and M.J.G.M. analysed the data and interpreted the results. M.T.P., M.J.G.M., J.E.V. and Y.C. wrote and reviewed the manuscript.

## Supplementary data

Supplementary data are available in the file Supplemetary.pdf.

## Data availability

All the data are publicly available as described in the Methods section. The simulation and analysis code to reproduce the results shown in this article are available from https://github.com/TamPham271299/DEJU

