## Supplementary materials for "Incorporating exon-exon junction reads enhances differential splicing detection"

December 16, 2024

### 1 Supplementary Figures

**Supplementary Figure S1**: Stacked barplots showing the average number of true (gray) and false (red) positive DEU genes at nominal 5% FDR detected by all 6 compared DEU analysis pipelines in different splicing pattern scenarios – ES, MXE, ASS, and IR across the combination of balanced/unbalanced library size and sample sizes (3,5 and 10 replicates per group). The observed FDR is shown over each bar. Results are averaged over 20 simulations with 75 bp paired-end read with 250 genuine DEU genes generated with a fold-change of 3 for each splicing pattern. Results of DEU/DEJU analyses are presented using both the gene-level Simes method and the $F$-test.

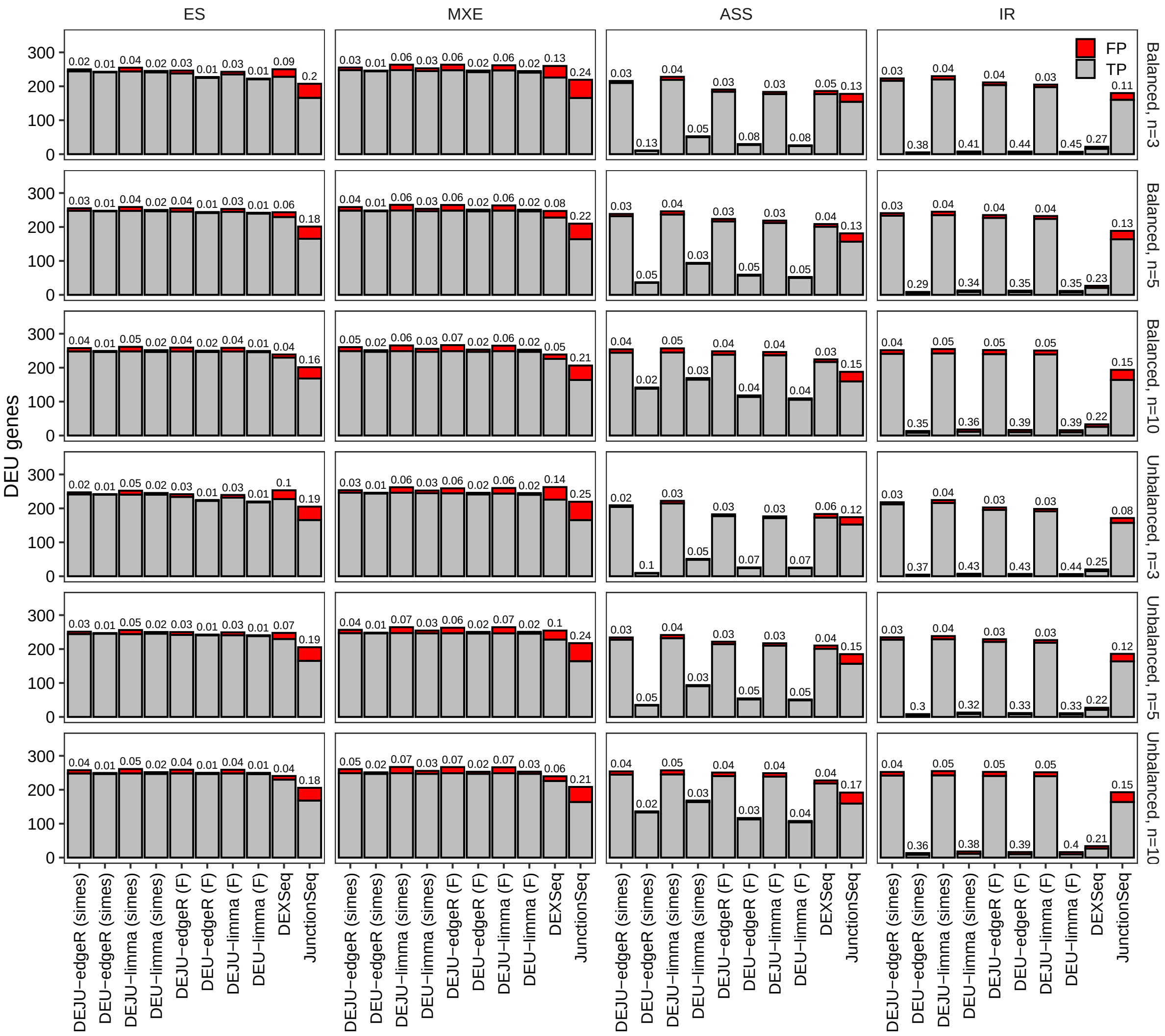

**Supplementary Figure S2**: Panels show the average number of false discoveries as a function of the number of chosen DEU genes across the combination of balanced/unbalanced library size and sample sizes (3,5 and 10 replicates per group). Results are averaged over 20 independent simulation runs with 75 bp paired-end reads. Results of DEU/DEJU analyses are presented using both the gene-level Simes method and the $F$-test.

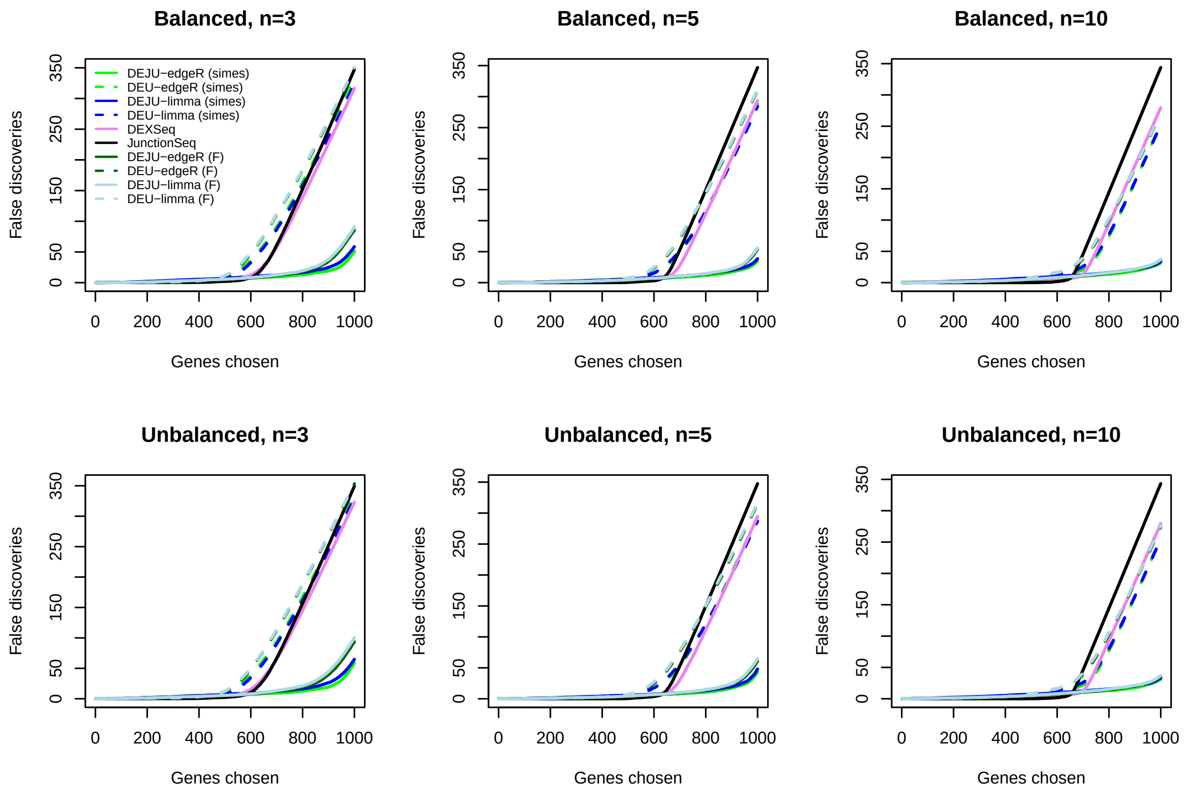

**Supplementary Figure S3**: Barplots with errorbars showing mean and standard error of the statistical sensitivity (or power) of the 6 compared pipelines in the DEU detection at nominal 5% FDR in 4 splicing pattern scenarios (ES, MXE, ASS, and IR) across the combination of balanced/unbalanced library sizes and sample sizes (3, 5, and 10 replicates per group). The averaged power is shown over each bar. Results are averaged over 20 simulations with 75 bp paired-end reads with 250 genuine DEU genes simulated with a fold-change of 3 for each splicing pattern. Results of DEU/DEJU analyses are presented using both the gene-level Simes method and the $F$-test.

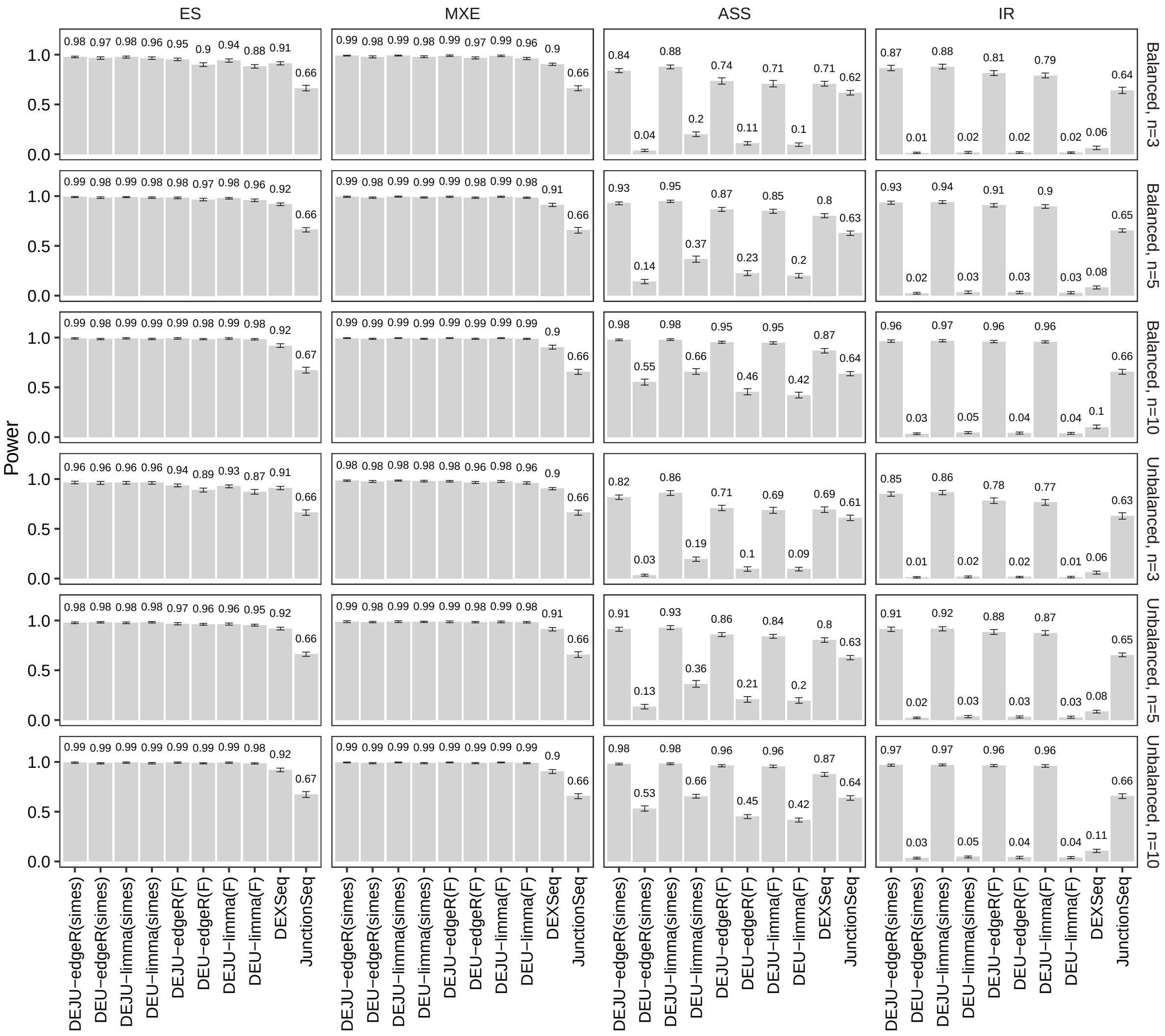

**Supplementary Figure S4**: Line plots showing the average and standard errors of the statistical power of 6 compared pipelines in the DEU/DEJU analyses with increasing numbers of replicates and balanced/unbalanced library sizes. Results of DEU/DEJU analyses are presented using both the gene-level Simes method and the $F$-test.

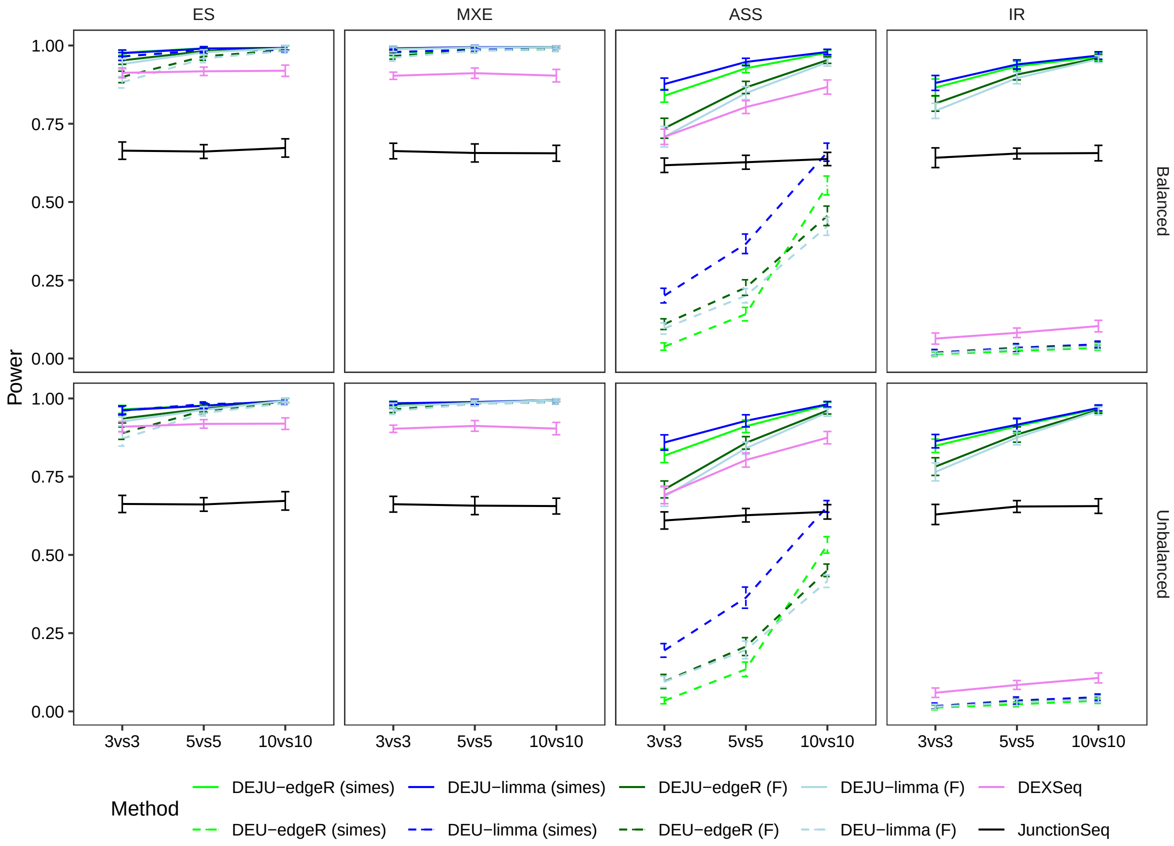

**Supplementary Figure S5**: Diagnostic plots of exon-junction read quantification by DEU-edgeR (A) and DEJU-edgeR (B).

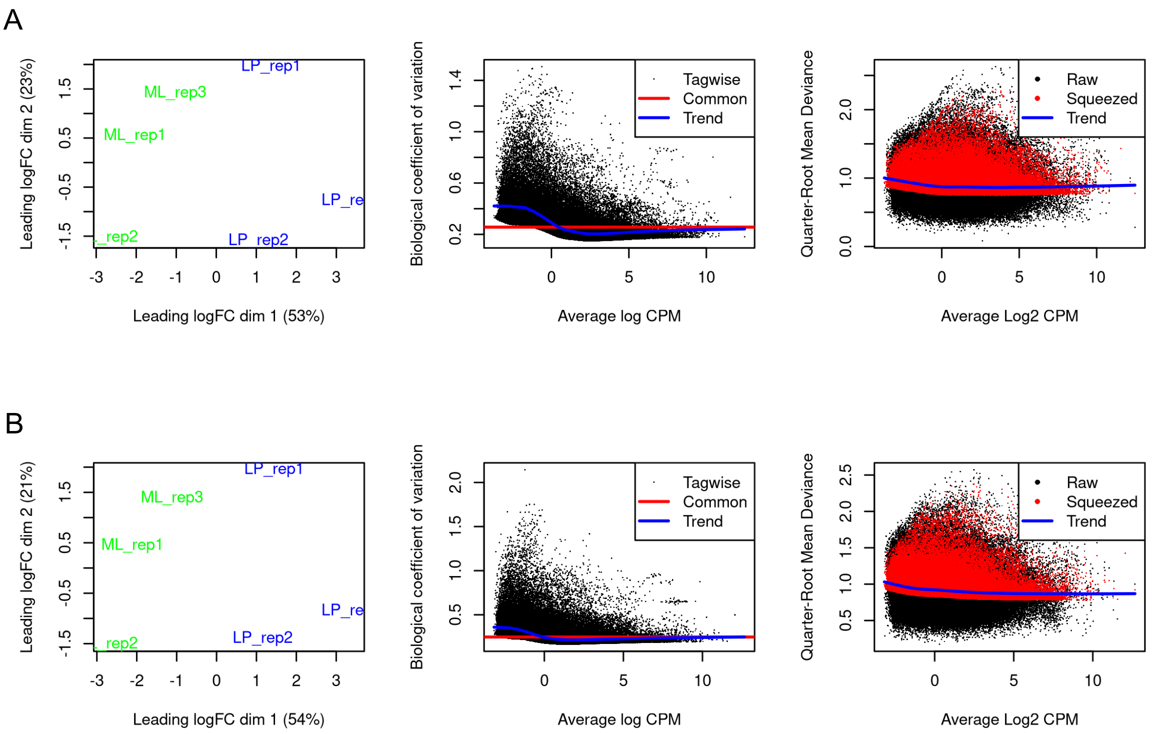

**Supplementary Figure S6**: (A) Illustrative examples of DEU genes detected by *DEJU-edgeR* not by *DEU-edgeR* using the gene-level Simes method or $F$-test): Myl6 (A, adjusted $P$-value = 9.4 x $10^{-6}$), Dusp16 (B, adjusted $P$-value = 1.7 x $10^{-3}$), Retreg1 (C, adjusted $P$-value = 0.02), and Mvb12a (D, adjusted $P$-value = 9.3 x $10^{-7}$). (Top panels) Schematic exon-junction plots showing the up-regulated (red), down-regulated (blue), and non-differential (black) usage of exons and junctions in LP versus ML samples; (Middle panels) a UCSC transcript annotation track of mm39; (Bottom panels) Sashimi plots alongside the coverage of RNA-seq reads of LP (blue) and ML (green) samples. Exon-exon junction read counts are shown in a box.

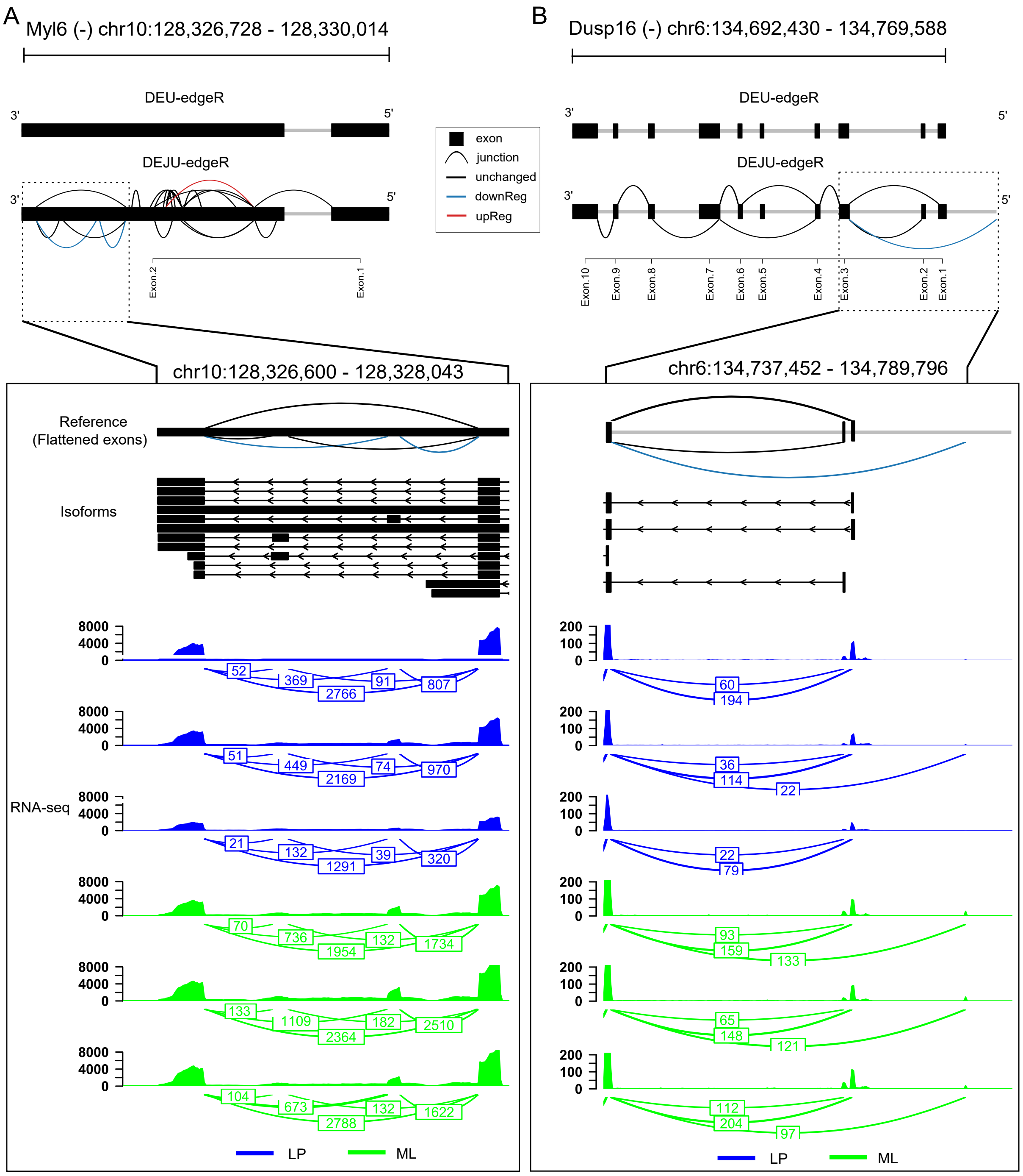

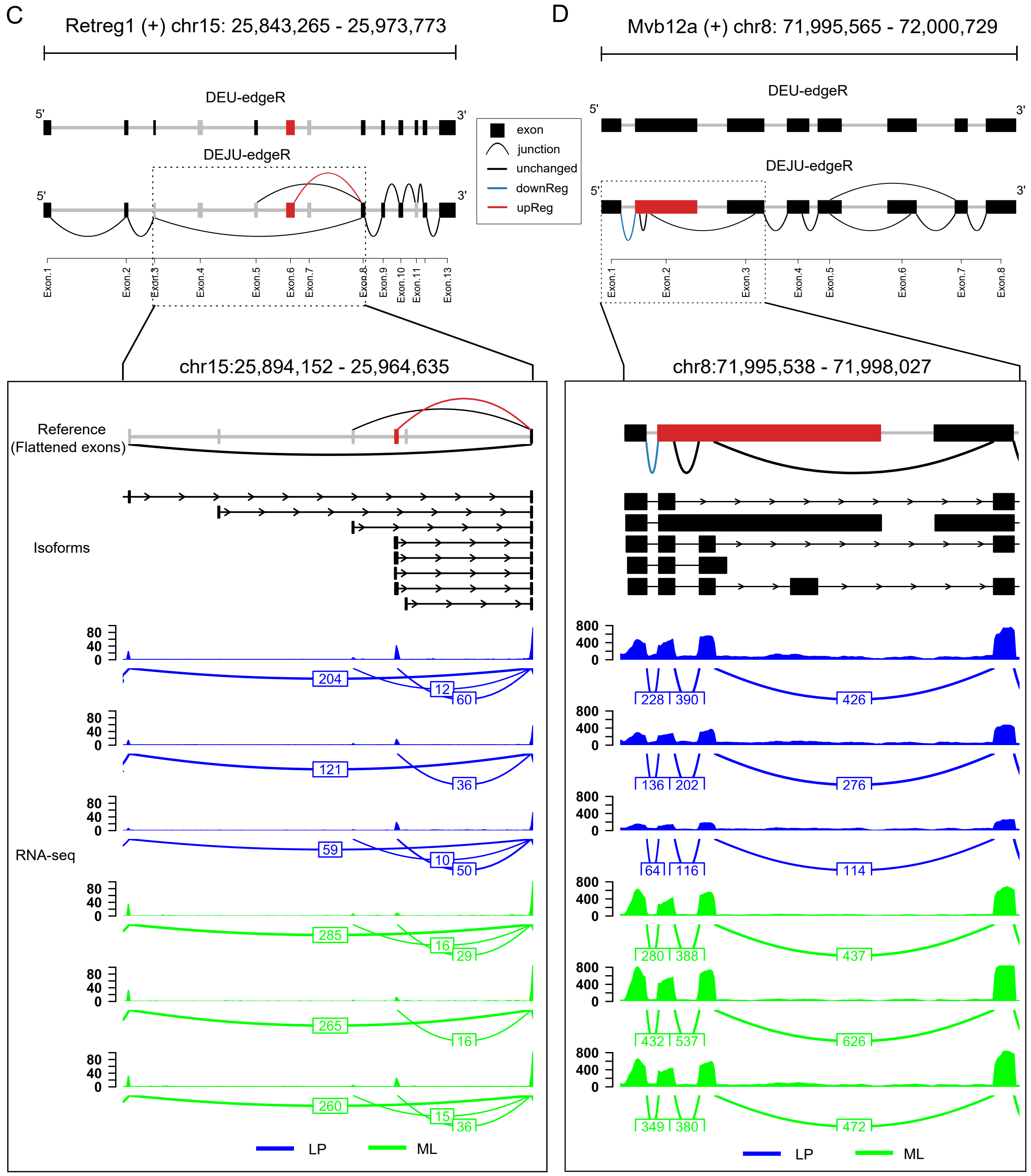
C
Retreg1 (+) chr15: 25,843,265 - 25,973,773
DEU-edgeR
DEJU-edgeR
5'
3'
Exon.1
Exon.2
Exon.3
Exon.4
Exon.5
Exon.6
Exon.7
Exon.8
Exon.9
Exon.10
Exon.11
Exon.13
chr15:25,894,152 - 25,964,635
Reference
(Flattened exons)
Isoforms
RNA-seq
204
12
60
121
36
59
10
50
285
16
29
265
16
260
15
36
LP
ML
exon
junction
unchanged
downReg
upReg
D
Mvb12a (+) chr8: 71,995,565 - 72,000,729
DEU-edgeR
DEJU-edgeR
Exon.1
Exon.2
Exon.3
Exon.4
Exon.5
Exon.6
Exon.7
Exon.8
chr8:71,995,538 - 71,998,027
228
390
426
136
202
276
64
116
114
280
388
437
432
537
626
349
380
472
LP
ML

**Supplementary Figure S7**: Upset plots illustrating the intersections of DEU gene sets detected in LP versus ML samples across six benchmarked pipelines. Results of DEU/DEJU analyses are presented using both the gene-level Simes method and the $F$-test.

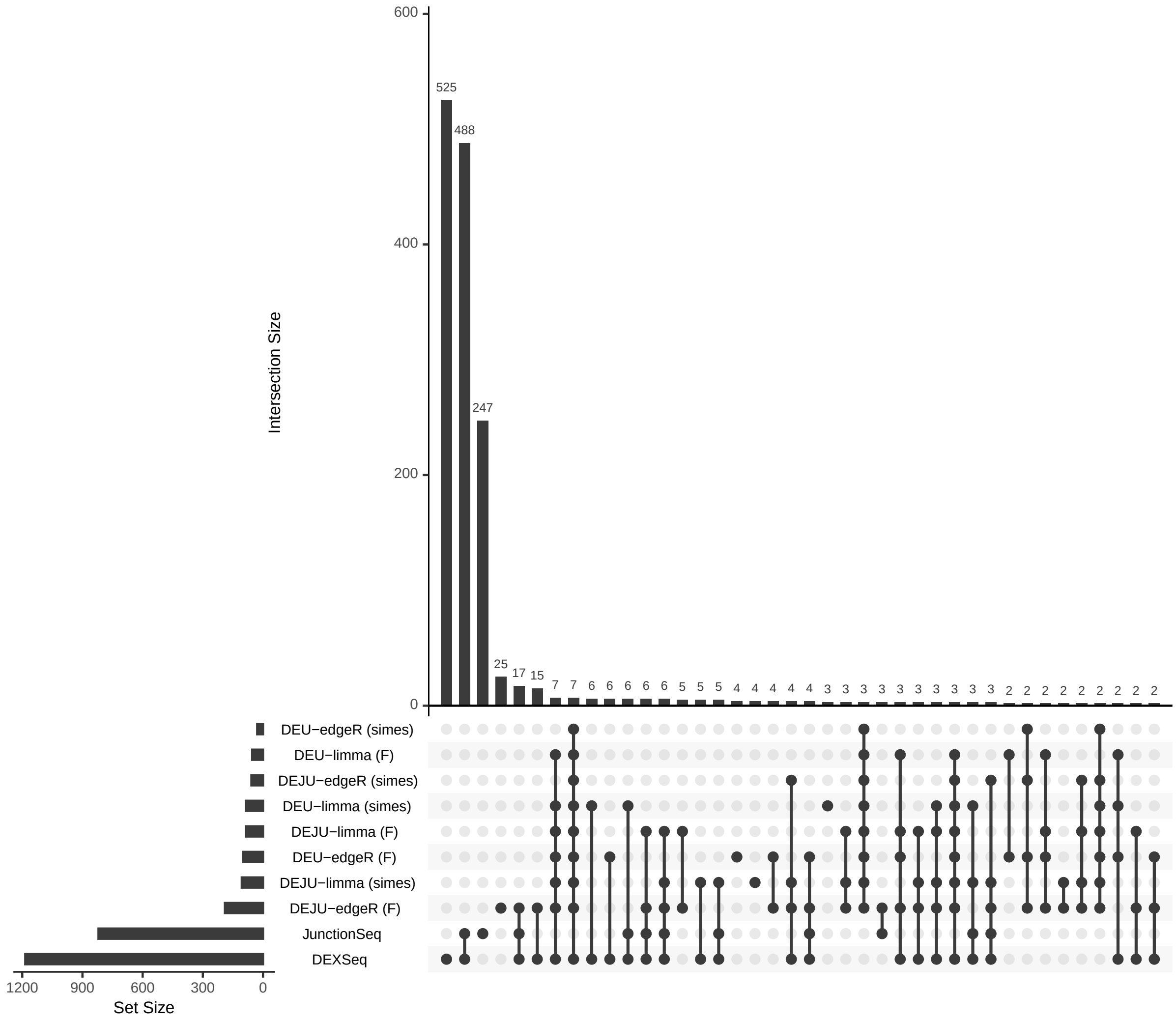

**Supplementary Figure S8**: Schematic exon-junction plots of representative DEU genes identified by *DEJU-limma* not by *DEU-limma* in LP compared to ML samples using the gene-level Simes method or *F*-test: Myl6 (B, adjusted *P*-value = 1.3 x $10^{-5}$), Dusp16 (C, adjusted *P*-value = 1.6 x $10^{-3}$), and Mvb12a (E, adjusted *P*-value = 6.7 x $10^{-7}$). Note that Fgfr1 (A, adjusted *P*-value = 0.06) and Retreg1 (D, adjusted *P*-value = 0.06) were not identified as DEU genes by *DEJU-limma*. Exons and splice junctions up-regulated, down-regulated, or non-differentially expressed in LP compared to ML samples are shown in red, blue, and black, respectively.

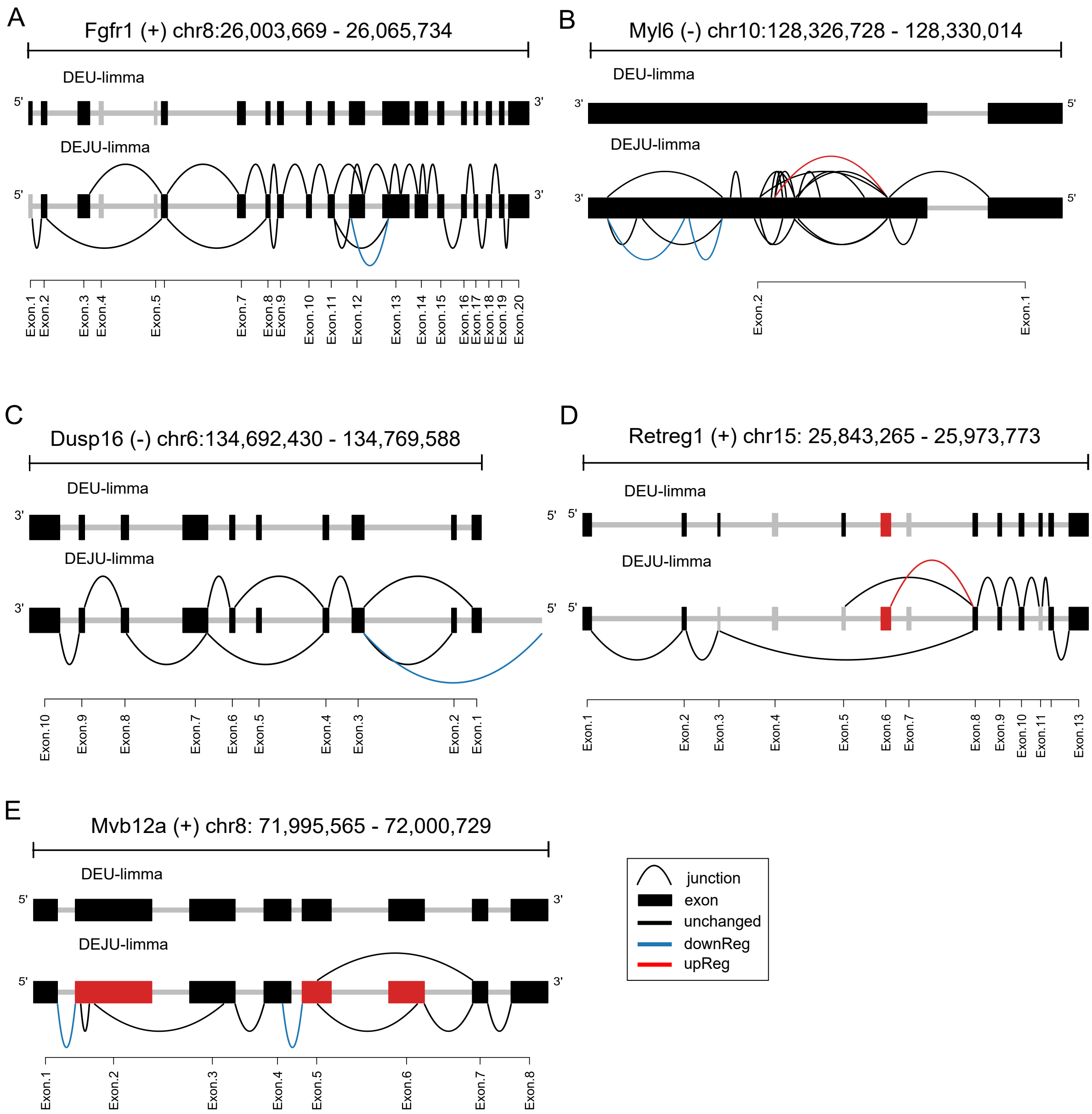

**Supplementary Figure S9**: Examples of DEU genes detected by *DEXSeq* in LP (red) compared to ML (blue) samples : Fgfr1 (A, adjusted $P$-value $< 2.2 \times 10^{-16}$), Myl6 (B, adjusted $P$-value $= 3.1 \times 10^{-6}$), Dusp16 (C, adjusted $P$-value $= 2.8 \times 10^{-8}$), Retreg1 (D, adjusted $P$-value $= 6.3 \times 10^{-4}$), and Mvb12a (E, adjusted $P$-value $= 1.2 \times 10^{-10}$). Exon bins hightlighted in pink are significant differentially used.

A

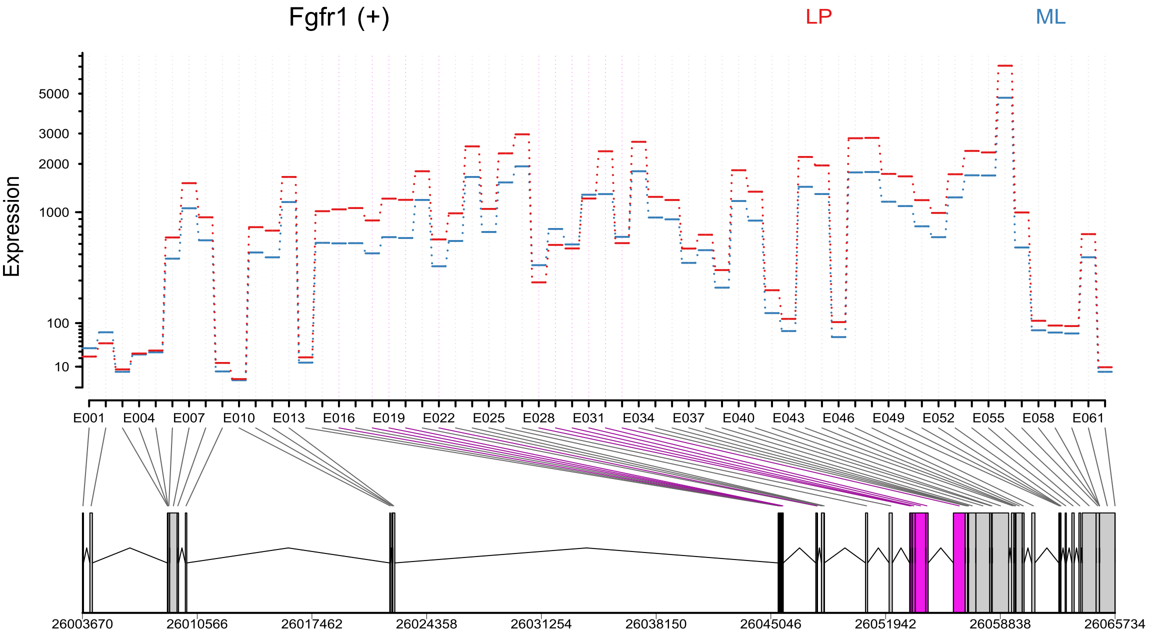

B

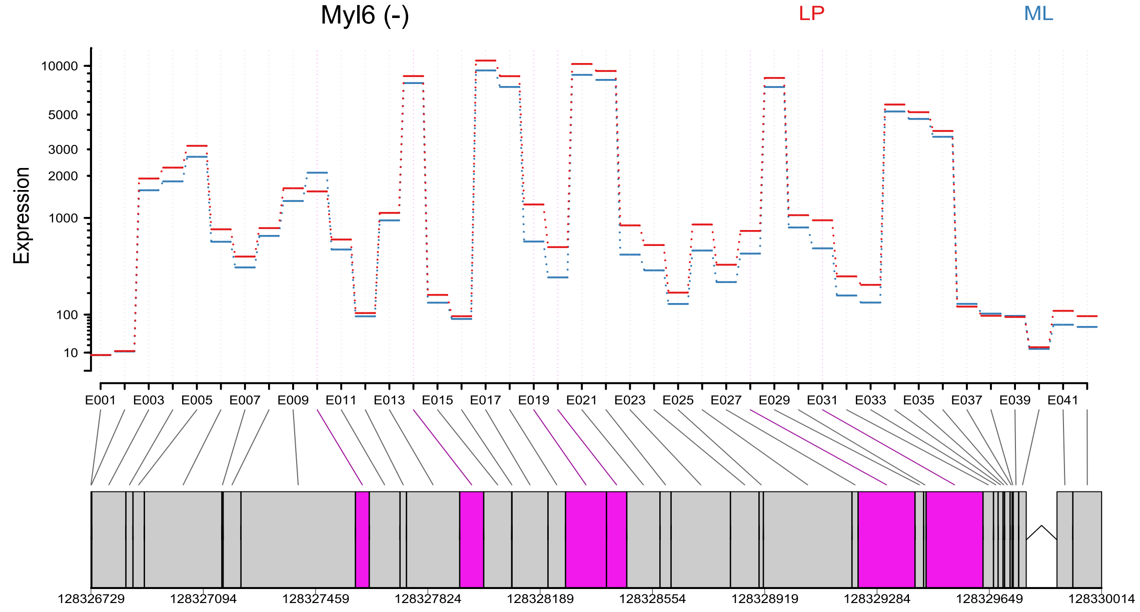

C

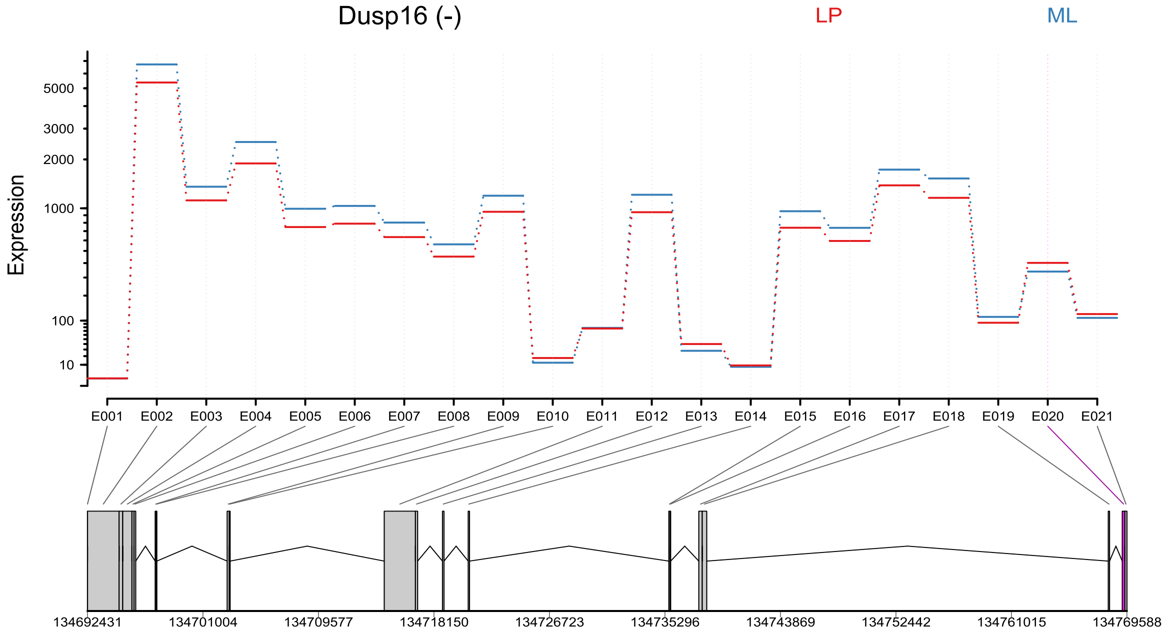

Dusp16 (-)
LP
ML
Expression
5000
3000
2000
1000
100
10
E001 E002 E003 E004 E005 E006 E007 E008 E009 E010 E011 E012 E013 E014 E015 E016 E017 E018 E019 E020 E021
134692431 134701004 134709577 134718150 134726723 134735296 134743869 134752442 134761015 134769588

D

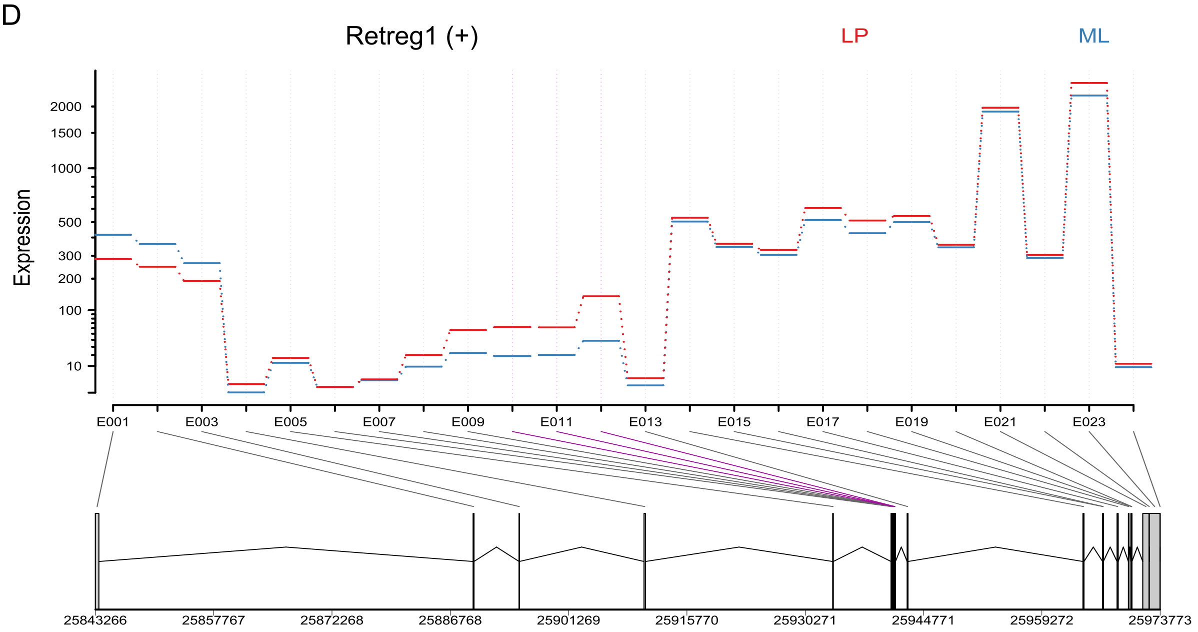

Retreg1 (+)
LP
ML
Expression
2000
1500
1000
500
300
200
100
10
E001 E003 E005 E007 E009 E011 E013 E015 E017 E019 E021 E023
25843266 25857767 25872268 25886768 25901269 25915770 25930271 25944771 25959272 25973773

E

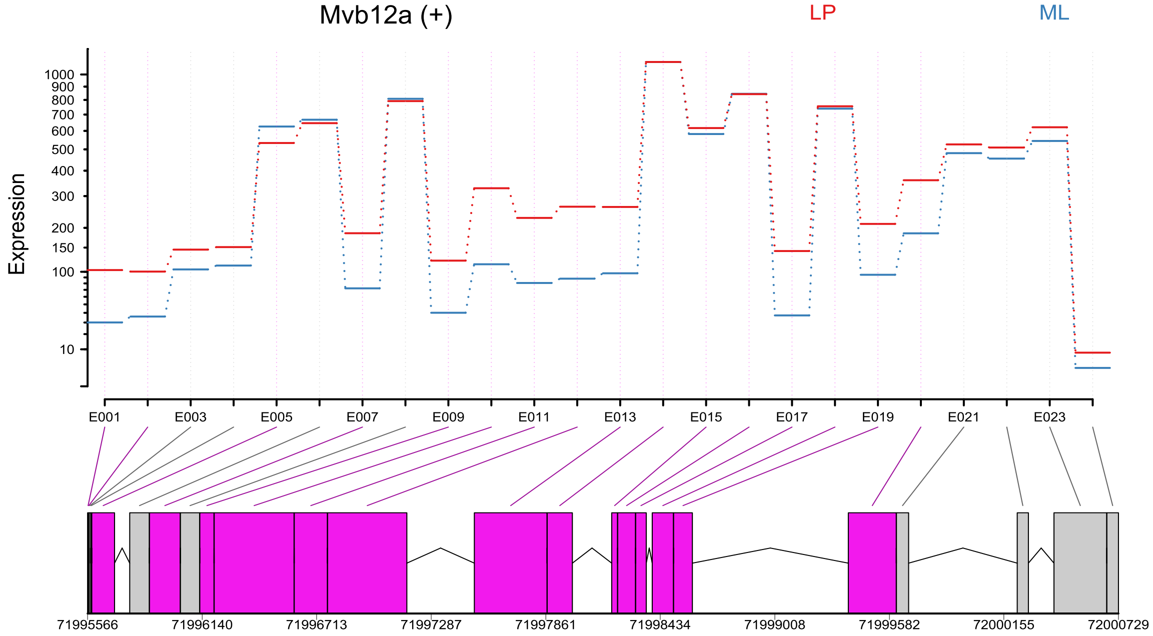
Mvb12a (+)
LP
ML
Expression
1000
900
800
700
600
500
400
300
200
150
100
10
E001
E003
E005
E007
E009
E011
E013
E015
E017
E019
E021
E023
71995566
71996140
71996713
71997287
71997861
71998434
71999008
71999582
72000155
72000729

**Supplementary Figure S10**: Examples of DEU genes detected by *JunctionSeq* in LP (blue) compared to ML (red) samples: Myl6 (A, adjusted $P$-value = 1.4 x $10^{-4}$), Dusp16 (B, adjusted $P$-value = 2.9 x $10^{-9}$), Retreg1 (C, adjusted $P$-value = 8.6 x $10^{-5}$), and Mvb12a (D, adjusted $P$-value = 8.4 x $10^{-10}$). Exon bins and splice junctions highlighted in pink are significant differentially used.

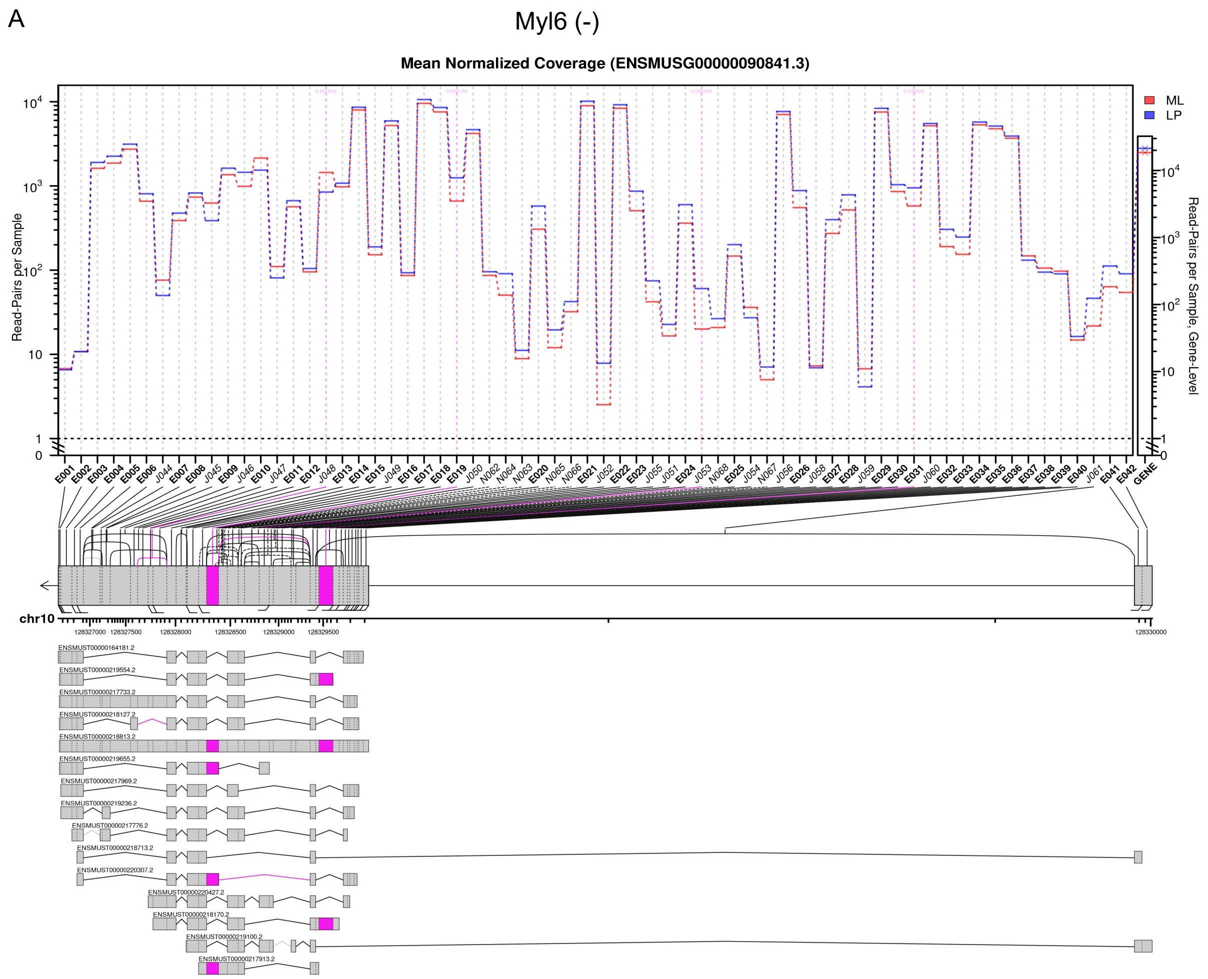

B

Dusp16 (-)

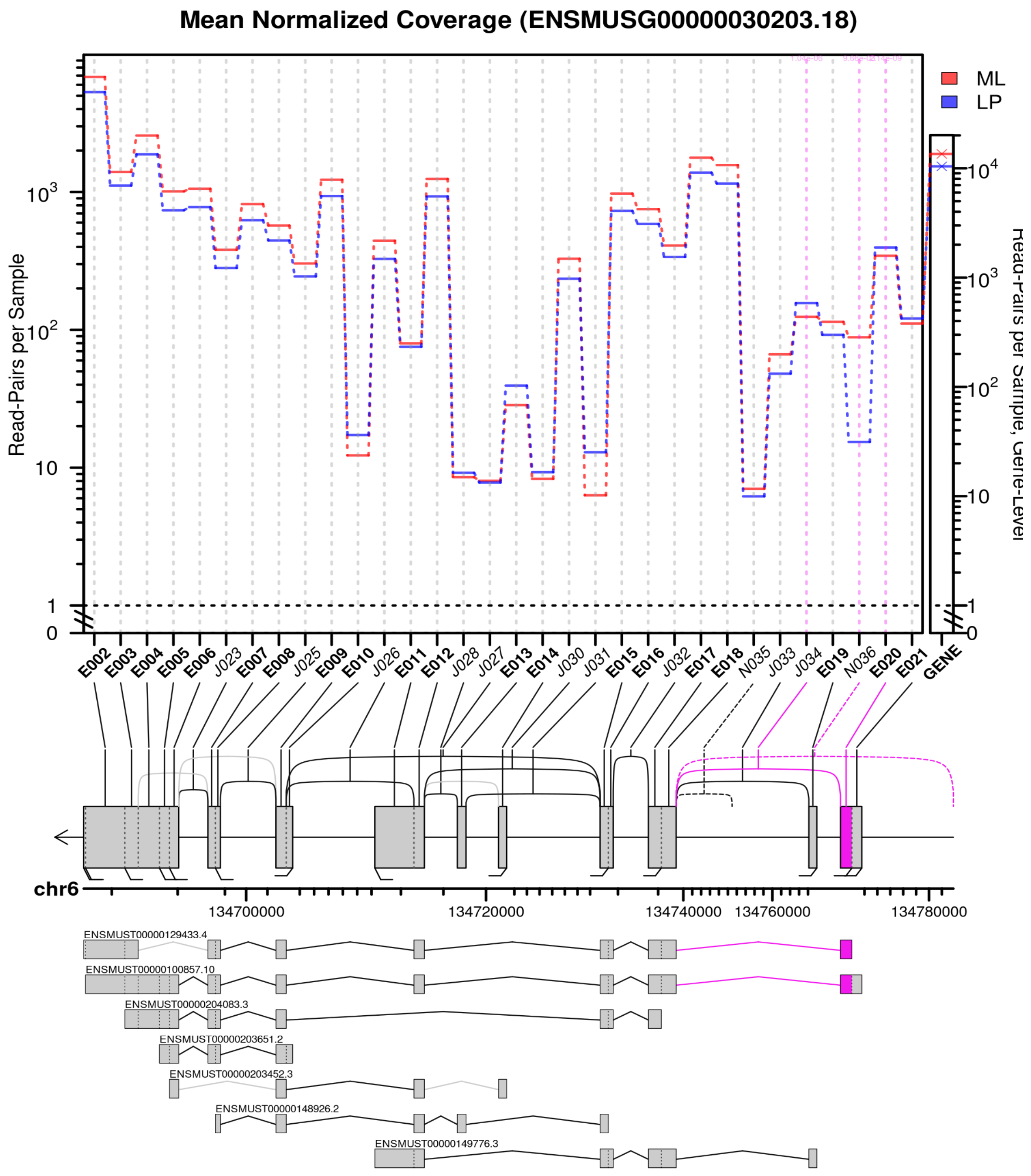

Mean Normalized Coverage (ENSMUSG00000030203.18)
Read-Pairs per Sample
Read-Pairs per Sample, Gene-Level
ML
LP
E002
E003
E004
E005
E006
J023
E007
E008
J025
E009
E010
J026
E011
E012
J028
J027
E013
E014
J030
J031
E015
E016
J032
E017
E018
N035
J033
J034
E019
N036
E020
E021
GENE
chr6
134700000
134720000
134740000
134760000
134780000
ENSMUST00000129433.4
ENSMUST00000100857.10
ENSMUST00000204083.3
ENSMUST00000203651.2
ENSMUST00000203452.3
ENSMUST00000148926.2
ENSMUST00000149776.3

C

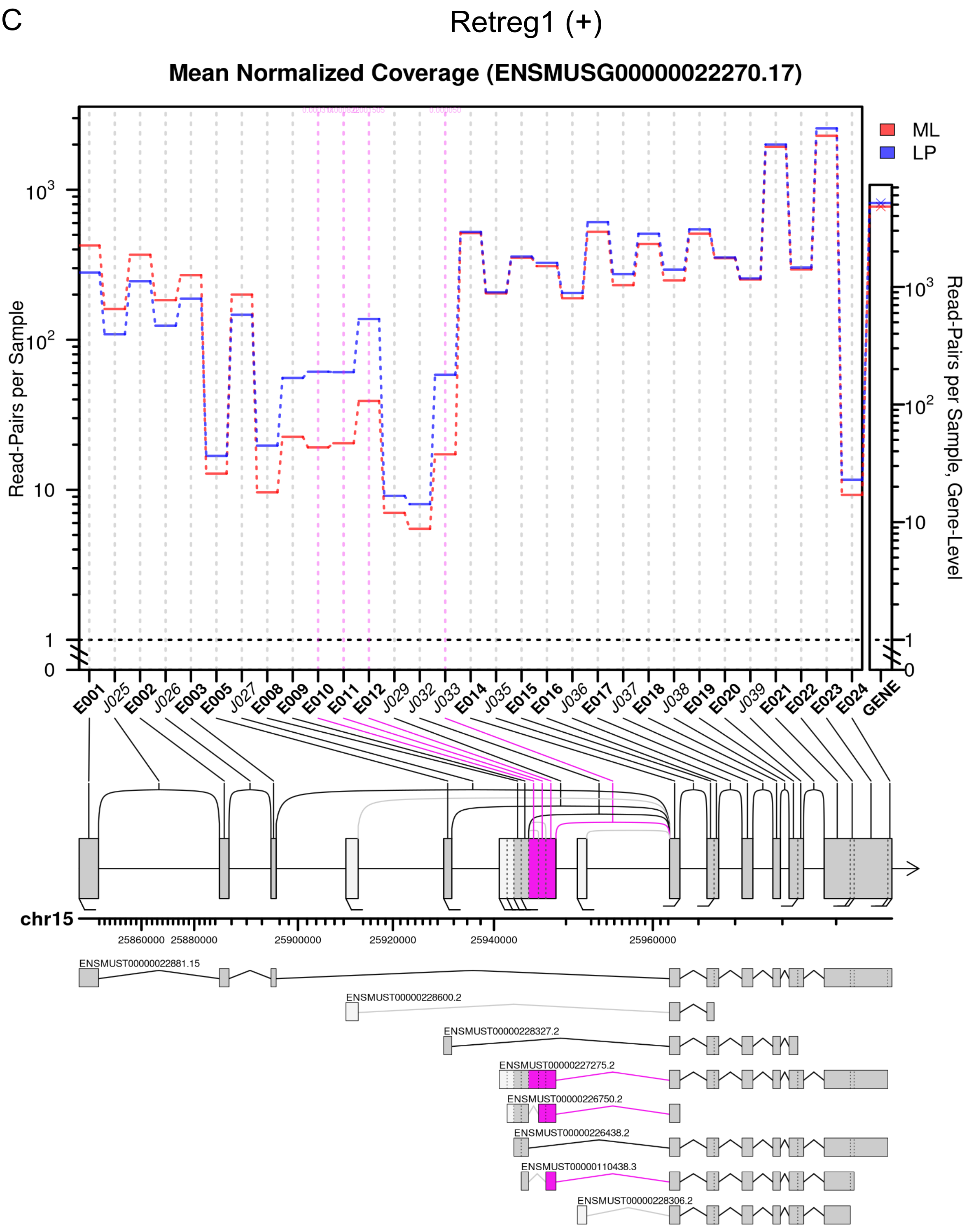

Retreg1 (+)
Mean Normalized Coverage (ENSMUSG00000022270.17)
Read-Pairs per Sample
Read-Pairs per Sample, Gene-Level
ML
LP
E001
J025
E002
J026
E003
E005
J027
E008
E009
E010
E011
E012
J029
J032
J033
E014
J035
E015
E016
J036
E017
J037
E018
J038
E019
E020
J039
E021
E022
E023
E024
GENE
chr15
25860000
25880000
25900000
25920000
25940000
25960000
ENSMUST00000022881.15
ENSMUST00000228600.2
ENSMUST00000228327.2
ENSMUST00000227275.2
ENSMUST00000226750.2
ENSMUST00000226438.2
ENSMUST00000110438.3
ENSMUST00000228306.2

D

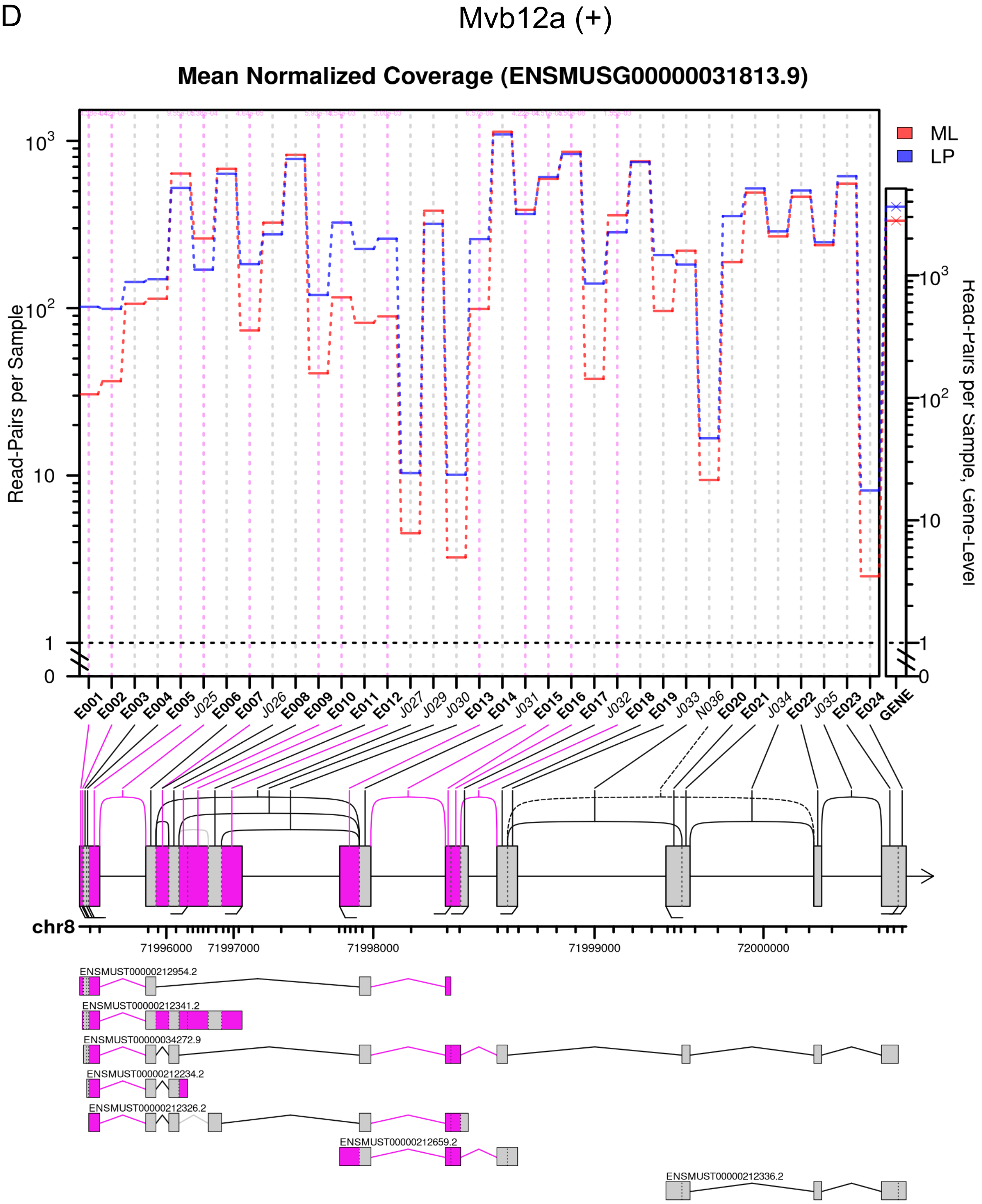

Mvb12a (+)
Mean Normalized Coverage (ENSMUSG00000031813.9)
ML
LP
Read-Pairs per Sample
Read-Pairs per Sample, Gene-Level
E001
E002
E003
E004
E005
J025
E006
E007
J026
E008
E009
E010
E011
E012
J027
J029
J030
E013
E014
J031
E015
E016
E017
J032
E018
E019
J033
N036
E020
E021
J034
E022
J035
E023
E024
GENE
chr8
71996000
71997000
71998000
71999000
72000000
ENSMUST00000212954.2
ENSMUST00000212341.2
ENSMUST00000034272.9
ENSMUST00000212234.2
ENSMUST00000212326.2
ENSMUST00000212659.2
ENSMUST00000212336.2

#### 2 Supplementary Tables

**Supplementary Table S1**: (Separate files) Summary of the average and standard error of FDR and statistical power in the DEU detection across six benchmarked pipelines, evaluated over 20 independent simulations in various scenarios with balanced/unbalanced library sizes and sample sizes (3, 5, and 10 replicates per group).

**Supplementary Table S2**: Table presents the walltime (min) and maximum memory usage (GB) of the DEU analysis pipelines tested in serial (1 core) and parallel (4 cores) modes. All tests were conducted on an Intel Xeon E5-2690 v4 server. Note: 1 GB = 1024 MB.

| **Libsizes** | **Samples per group** | **Method** | **Walltime (min)** | | **Memory (GB)** | |
|---|---|---|---|---|---|---|
| | | | **Parallel** | **Serial** | **Parallel** | **Serial** |
| Balanced | 3 | edgeR:diffSpliceDGE [a] | - | 0.47 | - | 0.64 |
| | | limma:diffSplice [a] | - | 0.33 | - | 0.55 |
| | | DEXSeq | 9.74 | 18.29 | 4.81 | 2.71 |
| | | JunctionSeq | 10.12 | 16.06 | 6.87 | 3.11 |
| | 5 | edgeR:diffSpliceDGE [a] | - | 0.64 | - | 0.63 |
| | | limma:diffSplice [a] | - | 0.42 | - | 0.63 |
| | | DEXSeq | 11.56 | 32.85 | 4.91 | 3.37 |
| | | JunctionSeq | 10.57 | 24.54 | 8.61 | 4.24 |
| | 10 | edgeR:diffSpliceDGE [a] | - | 0.71 | - | 0.80 |
| | | limma:diffSplice [a] | - | 0.41 | - | 0.75 |
| | | DEXSeq | 29.15 | 92.55 | 6.27 | 4.99 |
| | | JunctionSeq | 20.55 | 58.39 | 11.36 | 5.34 |
| Unbalanced | 3 | edgeR:diffSpliceDGE [a] | - | 0.50 | - | 0.58 |
| | | limma:diffSplice [a] | - | 0.32 | - | 0.58 |
| | | DEXSeq | 6.81 | 16.74 | 4.72 | 3.00 |
| | | JunctionSeq | 7.39 | 16.77 | 6.95 | 3.24 |
| | 5 | edgeR:diffSpliceDGE [a] | - | 0.55 | - | 0.58 |
| | | limma:diffSplice [a] | - | 0.35 | - | 0.59 |
| | | DEXSeq | 20.16 | 29.97 | 4.94 | 3.31 |
| | | JunctionSeq | 16.29 | 25.36 | 8.54 | 4.23 |
| | 10 | edgeR:diffSpliceDGE [a] | - | 0.93 | - | 0.74 |
| | | limma:diffSplice [a] | - | 0.60 | - | 0.72 |
| | | DEXSeq | 33.92 | 104.18 | 6.27 | 4.63 |
| | | JunctionSeq | 28.61 | 75.97 | 11.43 | 5.36 |

[a] The walltime and maximum memory usage for the *edgeR::diffSpliceDGE* and *limma::diffSplice* functions represent the total computational cost of both DEU and DEJU analyses. These functions do not support parallel processing.

**Supplementary Table S3**: Table shows the internal exon counts, junction counts, and library sizes of *DEU-edgeR* and *DEJU-edgeR* for three replicates of LP and ML samples in a case study by Milevskiy et al. (Milevskiy et al., 2023 – GSE227748).

| Group | Replicate | Internal exon reads | Junction reads | DEU-edgeR | DEJU-edgeR |
|---|---|---|---|---|---|
| | 1 | 38,519,524 | 35,306,601 | 102,722,690 | 73,824,068 |
| LP | 2 | 30,777,671 | 25,043,955 | 77,280,746 | 55,818,987 |
| | 3 | 20,625,053 | 16,710,546 | 52,589,308 | 37,333,995 |
| | 1 | 39,451,544 | 31,862,177 | 101,782,511 | 71,314,292 |
| ML | 2 | 46,352,497 | 36,445,354 | 117,546,848 | 82,799,073 |
| | 3 | 47,251,783 | 38,289,043 | 120,765,990 | 85,539,537 |

**Supplementary Table S4**: (Separate files) Lists of DEU genes of LP against ML samples exclusively detected by DEJU-edgeR using the gene-level Simes method and $F$-test.

**Supplementary Table S5**: Table shows the number of DEU genes in the comparison between LP and ML samples of 6 compared pipelines.

| | **Method** | **Gene-level test** | **Number of DEU detections** |
|---|---|---|---|
| LP vs ML | DEJU-edgeR | F | 190 |
| | | Simes method | 58 |
| | DEJU-limma | F | 85 |
| | | Simes method | 106 |
| | DEU-edgeR | F | 99 |
| | | Simes method | 29 |
| | DEU-limma | F | 54 |
| | | Simes method | 85 |
| | DEXSeq | | 1185 |
| | JunctionSeq | | 820 |

### 3 Supplementary Methods

#### 3.1 Reference genome

We retrieved the primary sequence FASTA file (GRCm39.primary_assembly.genome.fa.gz) and the comprehensive gene annotation GTF file (gencode.vM32.primary_assembly.annotation.gtf.gz) for the GRCm39 mouse genome (release M32) from the GENCODE database (`https://www.gencodegenes.org/`). The GTF file was used to generate comprehensive annotations for genomic features: flattened and merged exons and splice junctions. To create flattened and merged exon annotation, the *flattenGTF* function in the *Rsubread* Bioconductor package v2.16.1 (Liao *et al.*, 2019) with `GTF.featureType="exon", GTF.attrType="gene_id", method="merge"` options was employed to extract exonic regions from the GTF file and merge overlapping exons attributed to the same gene. The resultant exon annotation were saved in a standard annotation format (SAF) that contains five tab-separated columns: geneID, chromosomes, exon's start, exon's end, and strand. An example of a exon annotation SAF file is shown as below:

GeneID Chr Start End Strand
ENSMUSG00000000001.5 chr3 108014596 108016632 -
ENSMUSG00000000001.5 chr3 108016719 108016928 -
ENSMUSG00000000028.16 chr16 18599197 18599323 -
ENSMUSG00000000028.16 chr16 18600646 18600712 -
...

In addition, a splice junction annotation file, referred to hereafter as junction database, was generated from exonic regions at the transcript level. First, transcript-level exons were extracted from the GTF file. Splice junction coordinates were then defined as spanning from the last base of the upstream exon to the first base of the adjacent downstream exon. Duplicate junctions shared between transcript variants of the same gene were then merged to generate a set of unique junctions for each gene. The occurrence (or frequency) of junctions shared between different genes was also recorded. As a result, the junction database was saved in the tab-separated SAF format with extra columns including junctionID (Chr_Start_End) and junction's frequency. An example of a junction database is shown as below:

GeneID Chr Start End Strand JunctionID Frequency
ENSMUSG00000000001.5 chr3 108016632 108016719 - chr3_108016632_108016719 1
ENSMUSG00000000001.5 chr3 108016928 108019251 - chr3_108016928_108019251 1
ENSMUSG00000000028.16 chr16 18599323 18600646 - chr16_18599323_18600646 1
ENSMUSG00000000028.16 chr16 18600712 18603556 - chr16_18600712_18603556 1
...

#### 3.2 Simulation

##### Simulation of RNA-seq Data with Ground Truth

***Customizing Transcriptome With Designed Splicing Patterns.*** We randomly selected 5000 protein-coding genes with more than two exons to generate a customized transcriptome in which two transcripts per gene follow common alternative splicing patterns including Exon Skipping – ES, Mutually Exclusive Exon – MXE, Alternative 3'/5' Splice Site – ASS, and Intron Retention – IR in equal proportions (Figure **S11**).

With respect to the ES event, we randomly dropped one exon from each gene, generating two isoforms – a full-length isoform retaining all exons and a truncated isoform randomly omitting one exon. In the case of the MXE event, we randomly dropped one exon to generate one isoform and another exon to generate the second isoform, thereby creating two isoforms with mutually exclusive exons. For the ASS event, we randomly selected an exon and created a novel splice site inside the exon, either in the 3' or 5' direction. This process yields both a full-length and a truncated isoform with an alternative splice site. To mimic the IR event, we randomly selected two adjacent exons and fused them to create an intron-retained exon.

We converted flattened and merged exon annotation in SAF format into a transcript annotation in BED12 format with each row representing transcript's genomic intervals using the *bedtools* v2.31.1 (Quinlan and Hall, 2010) function *groupby*. We then used the *getfasta* function in *bedtools* to extract genomic sequences for each transcript interval from the FASTA file, resulting in a transcriptome FASTA file that exhibited designed splicing patterns. A schematic pipeline for generating transcript sequences in FASTA format for the ES splicing pattern is shown in Figure S12.

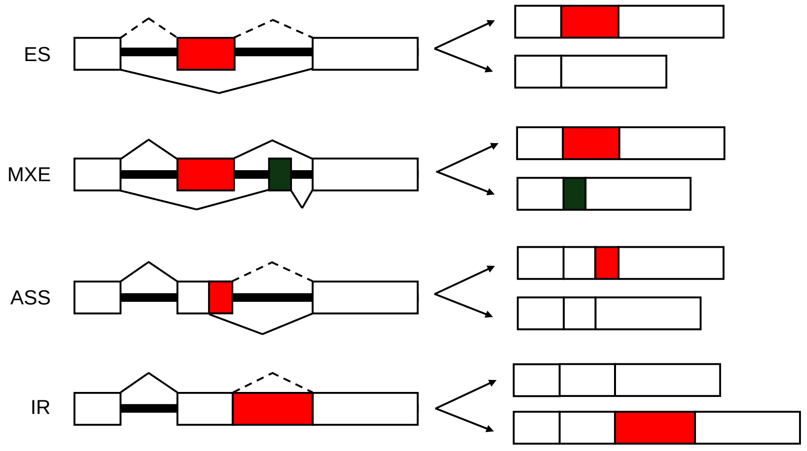

**Supplementary Figure S11**: Splicing patterns designed for the simulation datasets.

***Simulation of Ground Truth Data With and Without DEU genes.*** We created ground truth data through simulation of transcript-level expression under two scenarios, with and without any genuine DEU genes (null simulation) between two groups. Regarding simulation with DEU genes, we randomly selected 1000 out of 5000 genes as genuine DEU genes featuring four designed splicing patterns - ES, MXE, ASS, and IR - in equal proportions.

Baseline expression levels of transcripts were simulated following the Zipf's law (Furusawa and Kaneko, 2003). To simulate DEU genes, we first simulated the underlying abundance of one isoform for each gene following Zipf's law, subsequently simulating the underlying abundance of the second isoform by multiplying that of the first isoform with a random value ranging from 0.8 to 1.2 with a step of 0.01 in order to generate two isoforms with similar expression values. We then increased the abundance of one isoform in the first group and another isoform in the second group multiplicatively by a fold-change of 3. For the null simulation, the underlying abundance of all transcripts was simulated according to Zipf's law, with the same values across replicates of both groups.

Denote $\mu_{tij}$ the underlying abundance of transcript $t$ in the $j$-th replicate sample in the $i$-th group ($i$ = 1,2). Then $\mu_{tij}$ follows a gamma distribution with a shape $\phi_t^{-1}$ and a scale $\mu_{ti}\phi_t$, that is,

$$\mu_{tij} \sim \text{Gamma}(\phi_t^{-1},\ \mu_{ti}\phi_t), \qquad i = 1, 2$$

where $\phi_t$ follows the inverse Chi-square distribution, that is,

$$\phi_t \sim \text{Inv-}\chi^2(\phi_0, \text{df}_0),$$

with an underlying dispersion ($\phi_0$) of 0.05 and 10 degrees of freedom ($df_0$).

Resultant transcript-level expression values were in turn normalized relative to transcript length and the total number of expression values of each replicate, which were measured as transcripts-per-million (TPM). The normalized transcript expression data was finally employed to simulate RNA-seq data.

***Simulation of RNA-seq Reads.*** We used the *simReads* function in the Bioconductor package *Rsubread* to simulate RNA-seq reads in FASTQ format without sequencing errors for the customized transcriptome.

The simulation settings varied based on library size, which was either balanced at 50 million reads per sample or unbalanced, alternating between 25 million and 100 million reads across samples, and the number of biological replicates per group (3, 5, or 10). For each scenario, libraries consisting of 30 million paired-end reads, each 75 base pairs (bp) long, were generated for each replicate. Null simulations in which underlying transcript abundances remained consistent across replicates in both groups were also generated to assess FDR control.

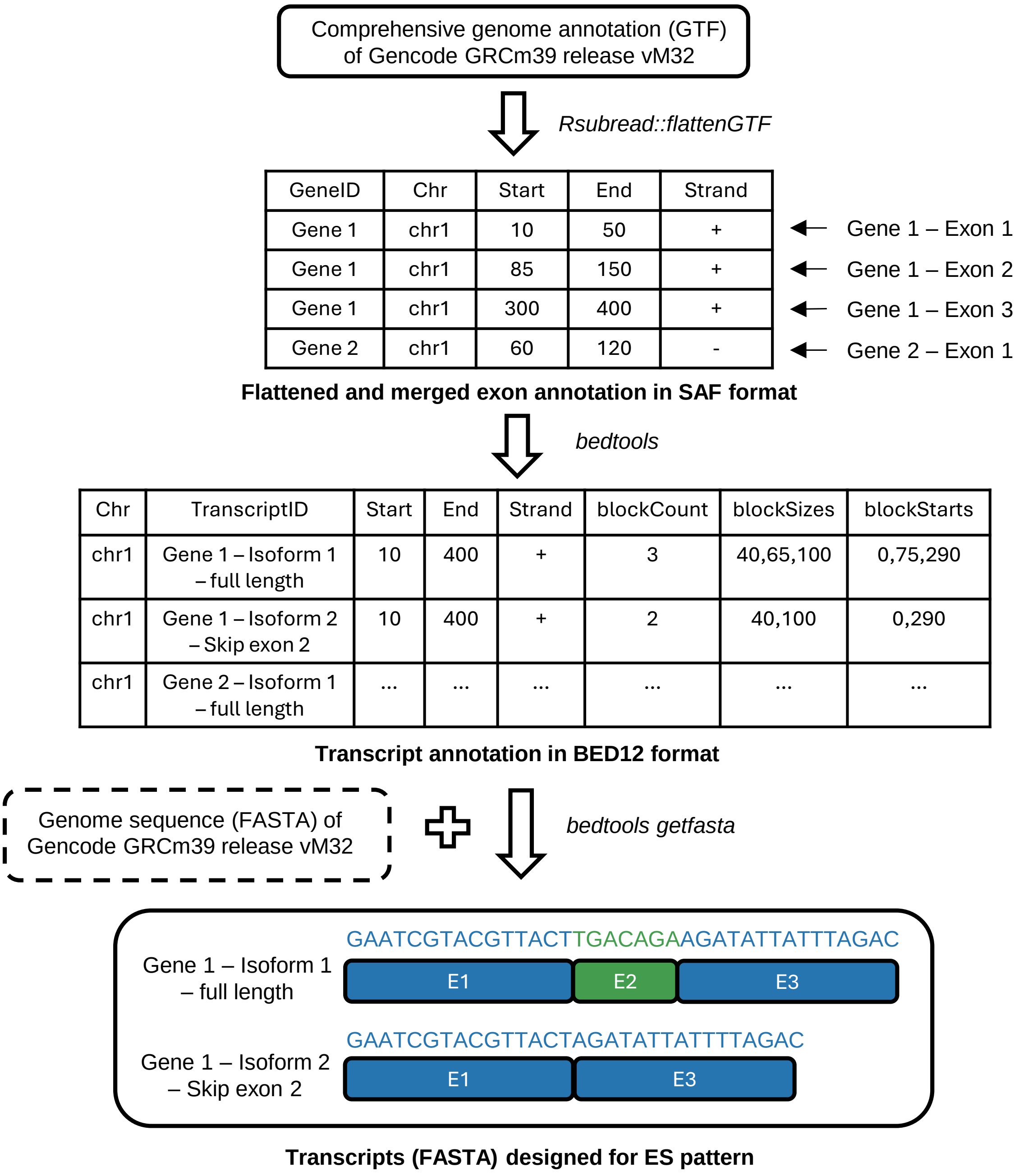

**Supplementary Figure S12**: A schematic workflow to generate transcripts (in FASTA format) customized for the ES pattern based on the mm39 reference genome annotation (GTF) and genome sequence (FASTA).

#### Analysis of RNA-seq Data

***Read Alignment.*** Processed RNA-seq reads were aligned using *STAR* v2.7.11b (Dobin *et al.*, 2013) in 2-pass mapping mode with a re-generated genome. This mode involves four main steps: genome indexing, 1st-pass mapping, optional filtering of splice junctions, and 2nd-pass mapping using a re-indexed genome generated using a set of junctions that passed filtering.

To index the genome, we used the genome annotation GTF file and reference genome sequence FASTA file with "`STAR --runMode genomeGenerate --genomeDir reidxGenome --genomeFastaFiles FASTA --sjdbGTFfile GTF --sjdbOverhang 74`" options. The parameter `sjdbOverhang` specifies the optimal splice junction overhang length, calculated as the maximum read length minus 1. For 75bp-long reads in the simulated FASTQ files, this value was set to 74.

In the first mapping pass, basic options were applied to generate alignment BAM files and junction count tables for each RNA-seq sample. Subsequently, junctions supported by fewer than two uniquely mapping reads (UMRs) across all RNA-seq samples were filtered out (Figure **S13**). This step ensures any junctions with very low counts in all samples of two groups should be discarded to improve the specificity and sensitivity in novel junction detection.

We then re-indexed the genome using the set of junctions that passed the filtering process (filtered.SJ.tab) with the command `"STAR --runMode genomeGenerate --genomeDir reidxGenome --genomeFastaFiles FASTA --sjdbOverhang 74 --sjdbFileChrStartEnd filtered.SJ.tab"`. The second mapping pass was performed with the additional parameter `--outFilterType BySJout`, which retains reads mapped to junctions passing junction-specific thresholds defined by `--outSJfilter*` parameters. By default, junction filtering settings mainly included the minimum overhang length on both sides for unannotated, annotated, canonical, and non-canonical junctions, which is 8, 1, 12, 30, respectively, and the minimum number of UMRs for non-canonical and canonical junctions, which is 3 and 1, respectively.

| | Group 1 | | | Group 2 | | | |
|---|---|---|---|---|---|---|---|
| | Rep 1 | Rep 2 | Rep3 | Rep 1 | Rep 2 | Rep 3 | |
| Junction 1 | 1 | 0 | 0 | 1 | 2 | 0 | → remove |
| Junction 2 | 1 | 2 | 0 | 10 | 7 | 12 | → keep |
| Junction 3 | 12 | 11 | 2 | 7 | 0 | 2 | → keep |

**Supplementary Figure S13**: An example of splice junctions filtered by the minimum number of UMRs (UMR cutoff of 2). Number of UMRs lower than the cutoff value are highlighted in grey.

##### Differential Exon and Junction Usage Analysis

***Exon-level, Internal Exon and Junction Read Quantification.*** Aligned reads were then quantified by the *Rsubread* function *featureCounts* (Liao *et al.*, 2014) using the flattened exon annotation SAF with the argument `useMetaFeatures=FALSE` to get exon-level read counts with doubled-counted exon-exon junction reads. An alternate counting strategy implemented in our proposed DEJU analysis pipeline classified exon-level reads into internal exon reads and exon-exon junction reads, which were summarized separately to get internal exon counts and junction counts. Particularly, internal exon counts were quantified by setting `nonSplitOnly=TRUE` to summarize only non-split reads, while junction counts were identified based on the "N" character in the CIGAR string, subsequently quantified by setting `juncCounts=TRUE`, with the reference genome sequence FASTA file used to summarize junction (or split) reads more precisely. Only uniquely mapping reads were counted in both methods using `minMQS=255` option.

***Incorporating Splice Junction Database.*** *featureCounts* was designed to automatically annotate each splice junction to a primary gene that overlaps with the highest number of splice sites compared to other genes, along with secondary genes that overlap with at least one of the two splice sites of that junction. In such cases where multiple genes overlap the same number of splice sites, *featureCounts* assigned the junction to the gene with the smallest leftmost base position as the primary gene. When different genes were nested within one another, *featureCounts* has shown to incorrectly assign annotated junctions as novel junctions of other genes, reducing the performance of downstream statistical analyses. To improve the accuracy of the junction annotation process, we used the junction database generated from the reference genome annotation GTF file as a reference to re-annotate junctions. Only unique, annotated junctions were considered. Figure **S14** shows an example of a unique, annotated junction with the junctionID of "chr1_20_100" that was re-assigned to the correct gene B based on the junction database.

***Read Filtering and Normalization.*** Lowly expressed exons and splice junctions were filtered by

| Primary Gene | Secondary Gene | Chr | Start | End | JunctionID |
|---|---|---|---|---|---|
| Gene A | Gene B | chr1 | 20 | 100 | chr1_20_100 |
| Gene C | Gene D | chr1 | 100 | 200 | chr1_100_200 |

Junction count table from *Rsubread::featureCounts*

+

| GeneID | Chr | Start | End | JunctionID | Frequency |
|---|---|---|---|---|---|
| Gene B | chr1 | 20 | 100 | chr1_20_100 | 1 |
| Gene C | chr1 | 100 | 200 | chr1_100_200 | 2 |
| Gene D | chr1 | 100 | 200 | chr1_100_200 | 2 |

Junction database

↓

| Primary Gene | Chr | Start | End | JunctionID |
|---|---|---|---|---|
| ~~GeneA~~ Gene B | chr1 | 20 | 100 | chr1_20_100 |
| Gene C | chr1 | 100 | 200 | chr1_100_200 |

Final junction count table

**Supplementary Figure S14**: Re-assignment of annotated junctions to the appropriate gene using the junction database.

the *edgeR* v4.0.16 (Robinson *et al.*, 2010) function *filterByExpr*. RNA compositions and library sizes were then normalized by *normLibSizes* function in the *edgeR* package, which provided the final exon-junction count matrix for the downstream DEJU analysis.

***Differential Exon and Junction Usage.*** For the DEU analysis, we performed differential usage of exons between two groups using the *edgeR* function *diffSpliceDGE*. For the DEJU analysis, differential exon and splice junction usage was carried out. Genes were considered to be DEU genes using the gene-level Simes method or $F$-test under the FDR control 0.05. Besides, significant differentially used exons and junctions between groups were identified using the exon-level test.

#### Execution of Other DEU Analysis Tools

***limma::diffSplice.*** We also conducted the DEU and DEJU analysis using *limma-voom* R/bioconductor package v3.58.1 (Ritchie *et al.*, 2015), following the same workflow as used with *edgeR*. The *diffSplice* function in *limma* was used to detect DEU genes using the gene-level Simes method or $F$-test. The exon-level t-test was used to identify the significant differentially used exons and junctions.

***DEXSeq.*** The quantification of exon reads and the DEU analysis were completed using the featureCounts-DEXSeq pipeline (`https://github.com/vivekbhr/Subread_to_DEXSeq`). First, an annotation GTF file that contains collapsed exon counting bins (DEXSeq.gtf) was prepared from the reference genome annotation GTF file using the Python script dexseq_prepare_annotation.py provided by *DEXSeq* package v1.48.0 (Anders *et al.*, 2012). After alignment BAM files were generated using *STAR* aligner, the *featureCounts* function of *subread* v2.0.6 was used to summarize reads mapping to collapsed exon bins using `"-p --countReadPairs -M -f -O -Q 255 -F 'GTF' -a DEXSeq.gtf"` options, which are the same as those specified in the read quantification step of the existing DEU analysis implemented in *egdeR* and *limma*. Exon counts were then used as inputs of the *DEXSeqDataSetFromFeatureCounts* function of *DEXSeq* that converted exon counts into DEXSeqDataSet object, followed by the *estimateSizeFactors*, *estimateDispersions*, *testForDEU*, *estimateExonFoldChanges* and *DEXSeqResults* functions to identify DEU genes. We also used the *perGeneQValue* function to obtain gene-level adjusted P-value. Aggregated genes that shared common exons were excluded from downstream analyses.

***JunctionSeq.*** Different from other benchmarked DEU analysis pipelines that analyzed exon and/or junction counts quantified by the *featureCounts*, *QoRTs* v1.3.0 (Hartley and Mullikin, 2015) was used to summarize exon-junction reads for *JunctionSeq* v1.16.0 (Hartley and Mullikin, 2016) to detect DEU genes. Particularly, *QoRTs* quantified reads mapping to collapsed exon counting bins and splice junctions using `--runFunctions writeKnownSplices,writeNovelSplices, writeSpliceExon` options. Exon-junction counts generated by *QoRTs* was then used as inputs of the *readJunctionSeqCounts* function of *JunctionSeq* which was in turn converted into a JunctionSeqCountSet object. Its size factors and dispersions was subsequently analyzed to fit the regression model and test for DEJU using a list of consecutive functions including *estimateJunctionSeqSizeFactors*, *estimateJunctionSeqDispersions*, *fitJunctionSeqDispersionFunction*, *testForDiffUsage* and *estimateEffectSizes* with default settings.

A summary of the RNA-seq read simulation, the read alignment and quantification, and the DEU/DEJU analyses is shown in Figure **S15**. Figure **S16** illustrates the differences between flattened and merged exon regions and exon bins. In our DEJU analyses implemented with *edgeR* and *limma*, both exon and junction reads were quantified, whereas existing DEU analyses only quantified exon reads. In contrast, DEXSeq quantified reads mapped to exon bins rather than merged exons. Extending beyond DEXSeq, JunctionSeq summarized reads mapped to both exon bins and splice junctions.

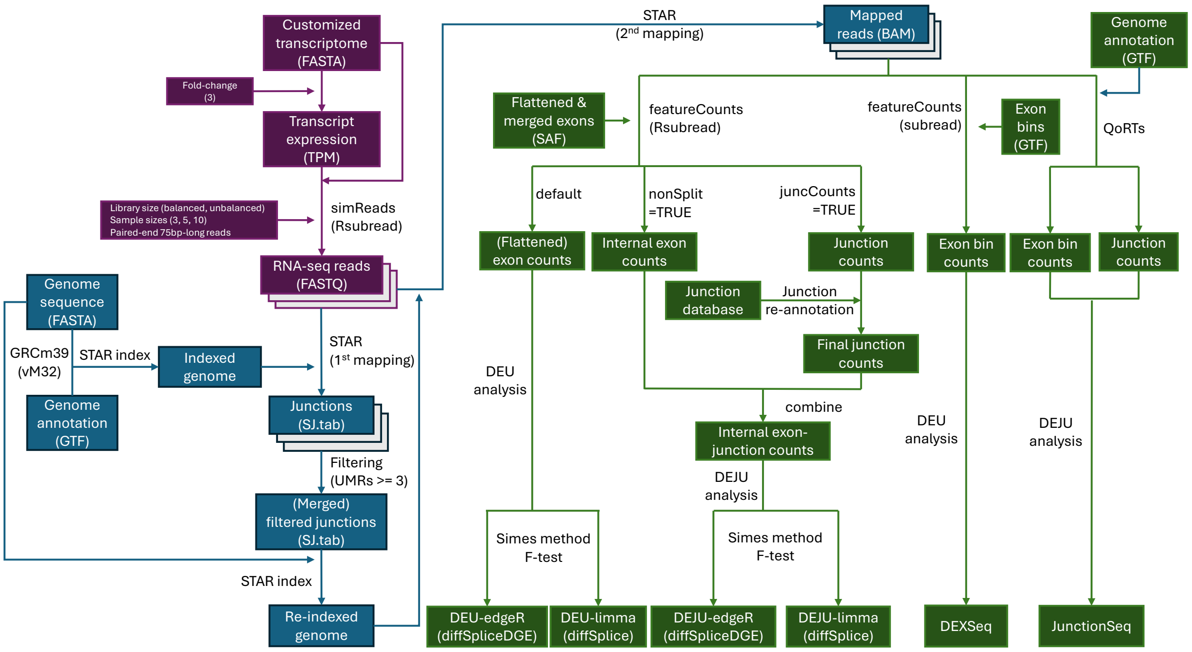

**Supplementary Figure S15**: A detailed workflow of the simulation of RNA-seq reads (purple), the read alignment (blue), and the quantification of exon-junction counts and DEU/DEJU analysis (green).

##### Benchmarking and Performance Assessment

We evaluated the performance of the DEJU analysis pipeline against the existing DEU pipeline and other popular tools including *DEXSeq* and *JunctionSeq* with respect to their statistical power (or sensitivity) in detecting DEU genes and the ability to control false discovery rate (FDR). In all DEU and DEJU analyses, significant results were identified under the FDR control of 0.05.

To calculate the FDR and power of each benchmarked method, we compared the DEU gene sets detected by each method with the reference set of simulated DEU genes. Let FP (false positive) denote DEU genes detected by the method but not in the reference DEU gene set; let FN (false negative) denote DEU genes in the reference set that were not detected; and let TP (true positive) denote the DEU genes in the reference set detected by the method.

The FDR was defined as

$$\text{FDR} = \frac{\text{FP}}{\text{TP} + \text{FP}}$$

$$\text{Precision} = 1 - \text{FDR}$$

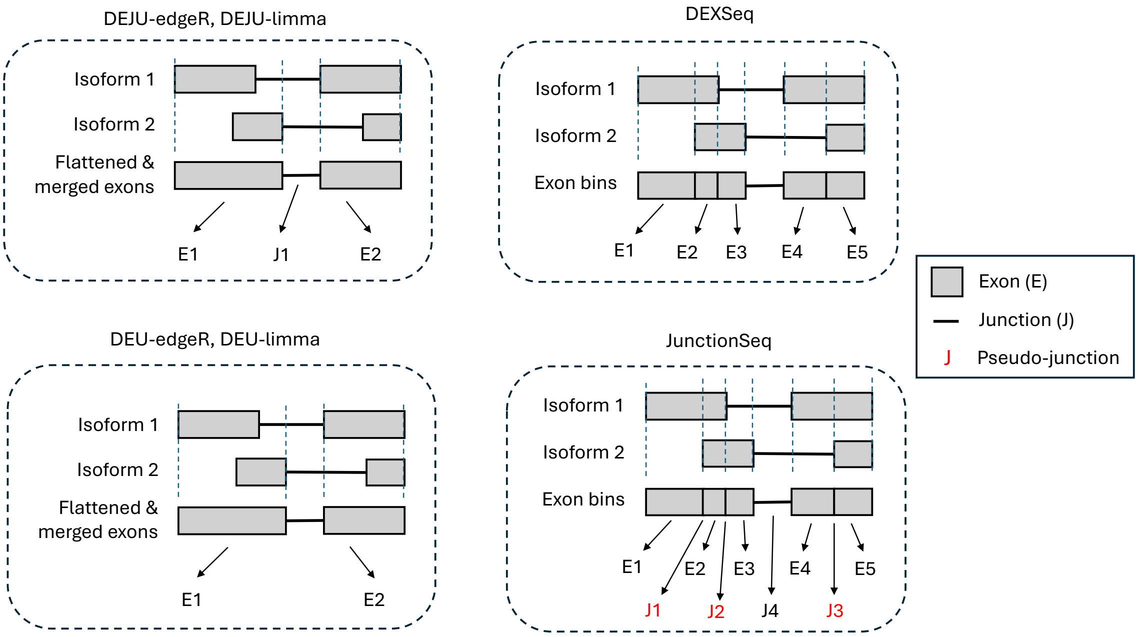

**Supplementary Figure S16**: Visualization of flattened and merged exon regions, as well as exon bins identified as features used in different DEU detection methods.

Correspondingly, the power was defined as

$$\text{Power (or Sensitivity)} = 1 - \frac{\text{FN}}{\text{TP} + \text{FN}}$$

#### 3.3 Case study

To evaluate the performance of our proposed DEJU analysis pipeline in the DEU detection, we analyzed RNA-seq FASTQ files of mouse mammary epithelial cells (MECs) including luminal progenitor (LP) and mature luminal (ML) from adult mice, which were sequenced using Illumina NextSeq 500 (GEO accesion number: GSE227748) (Milevskiy *et al.*, 2023). RNA-seq profiles of each cell type were sequenced in triplicate. We then used the similar workflow described in Figure **S15** with some modifications to improve the DEU detection from real RNA-seq experiments.

Since the main objective of the study is to evaluate the statistical analysis strategy of DEU detection methods, we simulated RNA-seq reads without sequencing errors. Therefore, quality control and read trimming were not essential for the simulated RNA-seq data. Yet, for real RNA-seq experiments, we performed these pre-processing steps to enhance overall alignment quality and ensure reliable downstream analyses. Quality control for the RNA-seq FASTQ files was performed using FASTQC v0.12.1 (`http://www.bioinformatics.babraham.ac.uk/projects/fastqc/`). Cutadapt v4.8 (Martin, 2011; Krueger, 2015) was then used with `"-q 30 --length 20 --paired --fastqc_args --nogroup"` options to remove adapter sequences of RNA-seq reads, trim low-quality ends from reads using a cutoff score of 20, and discard short reads with a minimum read length of 20bp.

For the read alignment step, since the maximum read length of FASTQ files is 81bp, the optimal splice junction overhang length was set to 80bp for indexing the reference genome. Besides, in the simulation process, due to the randomness in generating isoforms with designed splicing patterns, some of splice junctions were not real. Therefore, in order to detect artificial splice junctions for the simulation datasets, non-canonical junctions were considered. To improve the overall accuracy of the DEJU analysis for the real RNA-seq datasets, we excluded non-canonical junctions from our case-study analyses since most false positives of novel junctions are non-canonical. We used an extra option `--outFilterIntronMotifs RemoveNoncanonical` for the 2nd-pass mapping to exclude non-canonical junctions from the downstream statistical tests. In the DEU and DEJU analyses, genes without valid gene symbols were also discarded. Gene symbol and geneID were extracted from the GTF file.

**Visualization of DEU genes**

We used *Gviz* R/bioconductor package v1.46.1 (Hahne and Ivanek, 2016) to generate Sashimi plots to visualize RNA-seq read coverage and junction counts from BAM files. Other plots provided in the study were created using base R v4.3.3, *ggplot2* v3.5.0 (Wickham, 2016), *VennDiagram* v1.7.3 (Chen and Boutros, 2011), and *UpSetR* v1.4.0 (Conway *et al.*, 2017) R/bioconductor packages.
